# Septin-mediated coupling of protein import and division during chloroplast evolution

**DOI:** 10.64898/2026.03.28.715002

**Authors:** Samed Delic, Pamela Vetrano, Caroline S. Simon, David Su, Yezi Xiang, Shu-Zon Wu, Eva L. von der Heyde, Natsumi Tajima-Shirasaki, Sheng-An Chen, Carla Brillada, Armin Hallmann, Magdalena Bezanilla, Niccolo Banterle, Gautam Dey, Silvia Ramundo, Masayuki Onishi

## Abstract

Chloroplast biogenesis depends on both protein import and organelle division, yet how their coordination emerged during evolution remains unclear. Here, we show that the single septin SEP1 links these pathways in the green alga *Chlamydomonas reinhardtii*. SEP1 forms a filamentous network on the chloroplast envelope during interphase and reorganizes into a ring at the chloroplast division site during cytokinesis. Loss of SEP1 selectively impairs import of chloroplast-division proteins and causes mispositioning of the division ring, without impairing bulk chloroplast protein import. SEP1 physically associates with outer-envelope TOC GTPases through evolutionarily related GTPase domains. Phylogenetic analysis places TOC GTPases within an algal septin-derived clade, and heterologous expression of SEP1 in land plants, in which septins are absent, shows conservation of its chloroplast targeting and TOC binding. Together, these findings identify septins as coordinators of plastid protein import and division and suggest that this coupling emerged early in chloroplast evolution.

## INTRODUCTION

Chloroplasts originated from the primary endosymbiosis of a cyanobacterium by a heterotrophic eukaryote more than one billion years ago^1–3^. The emergence of photosynthetic eukaryotes that harbor this organelle dramatically increased atmospheric O_2_, thereby transforming life on Earth ^4,5^. During this transition, the eukaryotic host must have assumed responsibility for maintaining and replicating this new intracellular resident. For this ancient organism, monoplastidy was likely selected for to secure proper vertical inheritance, a trait seen in early diverging lineages of Archaeplastida ^6,7^. While exact mechanisms of how the host cell gained control over this division process remain unclear, many cyanobacterial genes (including those related to division) were transferred to the host nuclear genome, creating a dependency on nuclear-encoded proteins for chloroplast function ^8,9^, a process called endosymbiotic gene transfer (EGT). To restore the function of these transferred genes, eukaryotic cells evolved dedicated import machinery to deliver their protein products back into the organelle ^10–12^. In parallel, permanent organelle inheritance required that chloroplast division become coordinated with host cell division ^13,14^. Although these three processes are foundational to plastid evolution, how they may have become mechanistically integrated throughout the bacteria-to-organelle transition remains unclear. Monoplastidic algae such as *Chlamydomonas reinhardtii* are powerful model systems for dissecting how early eukaryotes evolved control over endosymbiont division, because cell division and chloroplast division are spatiotemporally coupled, and the system is genetically tractable.

Chloroplast division depends on a set of nuclear-encoded proteins that must be imported into the organelle. In Viridiplantae, this includes two FtsZ homologs, FTSZ1 and FTSZ2, which co-assemble into a heteropolymer known as the FTSZ-ring ^15–17^. This ring forms at the chloroplast midzone and is anchored to the inner envelope by ARC6 ^18–20^. Both FTSZ and ARC6 are derived from the ancestral cyanobacterium and were transferred to the host nuclear genome through EGT ^19,21^. As a result, these proteins are synthesized in the cytosol and imported into the chloroplast via the chloroplast-translocon system. Typical chloroplast-targeted proteins contain N-terminal targeting peptides ^22–24^. The TOC (Translocon of the Outer envelope of the Chloroplasts) complex recognizes these targeting peptides via conserved GTPase receptors such as TOC33 and TOC159, initiating translocation. In *Arabidopsis thaliana,* both TOC33- and TOC159-family proteins have undergone recent gene duplications. The resulting paralogs show overlapping yet distinct substrate preferences, suggesting a degree of specialization in precursor recognition ^25,26^. Precursor proteins targeted for the stroma are then passed through the TIC (Translocon of the Inner envelope of the Chloroplasts) complex and subsequently processed to remove the targeting peptides. The evolutionary origins of these translocon GTPases remains a mystery, although some work has speculated on the characteristics of their ancestral form^27^. Reumann et al. have stated ^28^, “It thus appears that the receptor of the TOC complex derive from an ancient eukaryotic protein that was encoded in the nucleus prior to branching of the eukaryotic lineage into plants and mammals.”

The import machinery is often assumed to function constitutively, acting as a static channel for protein delivery. However, some evidence suggests that the translocon is developmentally regulated to modulate chloroplast biogenesis and chloroplast division ^29^. Gain-of-function mutations in the TIC236 translocase suppress chloroplast division defects in *crumpled leaf (crl)* mutants of *A. thaliana*, raising the possibility that protein import and division are functionally connected ^30^. In another study, chloroplast division defects were observed in maize *defective kernel 5 (dek5)* mutants, which correlated with decreased levels of chloroplast translocon components ^31^. These findings highlight the importance of efficient translocon activity for importing stromal division proteins. However, whether translocon composition or function is modulated to facilitate chloroplast division remains an open question.

Septins are a conserved family of GTP-binding proteins most well studied in yeasts and animals (opisthokonts), where subunits encoded by multiple genes form short, heterooligomeric, and palindromic protomers in defined stoichiometries ^32,33^. These protomers then polymerize end-on-end on membrane surfaces into filaments, and further reassemble into higher-order structures such as rings at regions of micron-scale curvature ^34–37^. These septin structures are found on the plasma membrane, at mitochondrial fission sites ^38–40^, on phagocytic vesicles ^41^, and on the surface of intracellular bacterial pathogens ^42^. These various localizations have been almost exclusively described in opisthokont systems, with few exceptions ^43–45^. Recent phylogenetic studies conducted pan-eukaryotic searches for septins and found their conservation in both Chromista and Archaeplastida ^46–48^, indicating a deep evolutionary origin for this protein family at the level of the last eukaryotic common ancestor (LECA) ^48^. Furthermore, many non-opisthokont species encode a single septin in their genomes, suggesting that LECA had just one septin that may have functioned as a monomer or homo-oligomer. Despite their broad phylogenetic distribution, little is known about the cellular functions of septins outside opisthokonts; therefore, we currently lack an understanding of the ancestral septin functions.

Here, we characterize the single septin in *Chlamydomonas,* SEP1, and uncover its role in coordinating chloroplast protein import and division. SEP1 localizes to the chloroplast envelope, forms filaments during interphase, and assembles into a ring at the chloroplast division site. SEP1 physically interacts with outer membrane TOC GTPase components. Loss of SEP1 impairs both the positioning and import of stromal division proteins, despite normal TOC protein levels and envelope targeting. Phylogenetic analysis reveals that TOC GTPases are derived from an ancestral algal septin subfamily, and SEP1 retains the ability to localize and bind translocon proteins across diverse species, including land plants in which septins have been lost. These findings reveal a previously unrecognized mechanism linking translocon organization to organelle division and suggest that septins may have coordinated these processes during early plastid evolution.

## RESULTS

### Single *Chlamydomonas* septin SEP1 localizes to the chloroplast outer membrane and forms a ring during chloroplast division

While septins have long been studied in animal and fungal model systems, knowledge on their functions in other eukaryotic lineages is limited: In the ciliate *Tetrahymena thermophila*, septins localize to mitochondria ^44^, and in two green algae, *Nannochloris bacillaris* and *Marvania geminata*, septins have been localized to the division plane by immunostaining of fixed cells^43^*. Chlamydomonas* is one of many algae whose genome encodes a single septin gene ^45,48,49^. This single septin, SEP1, contains the conserved septin G domain core (G1, G3, and G4 motifs) and the septin unique element (SUE), and encodes a C-terminal amphipathic helix (AH), a feature implicated in septin-membrane interactions ^48,50^. SEP1 has been reported to form a homodimer through its G-interface *in vitro* ^45^. However, its organization and function within the cell remain unknown (Fig. 1A). To determine the subcellular localization of the single *Chlamydomonas* septin, we endogenously tagged SEP1 with a C-terminal fluorescent marker (SEP1-NG-3FLAG) using CRISPR-based genome editing (Supplementary Fig. 1A). Tagged clones showed loss of the wild-type SEP1 allele by PCR and expected band-shifting for the tagged cassette (Supplementary Fig. 1B). Multiple independent clones displayed consistent localization of fluorescent protein expression and co-segregation of the SEP1-NG cassette with hygromycin resistance (Supplementary Fig. 1C-D), indicating single-locus insertion. Live-cell imaging revealed that SEP1 formed small enrichments at the chloroplast envelope, with enhanced signal at discrete foci along the organelle surface (Fig. 1B). Co-expression with the outer envelope marker LCIA-SC confirmed its colocalization with SEP1 (Fig. 1B).

**Figure 1.**
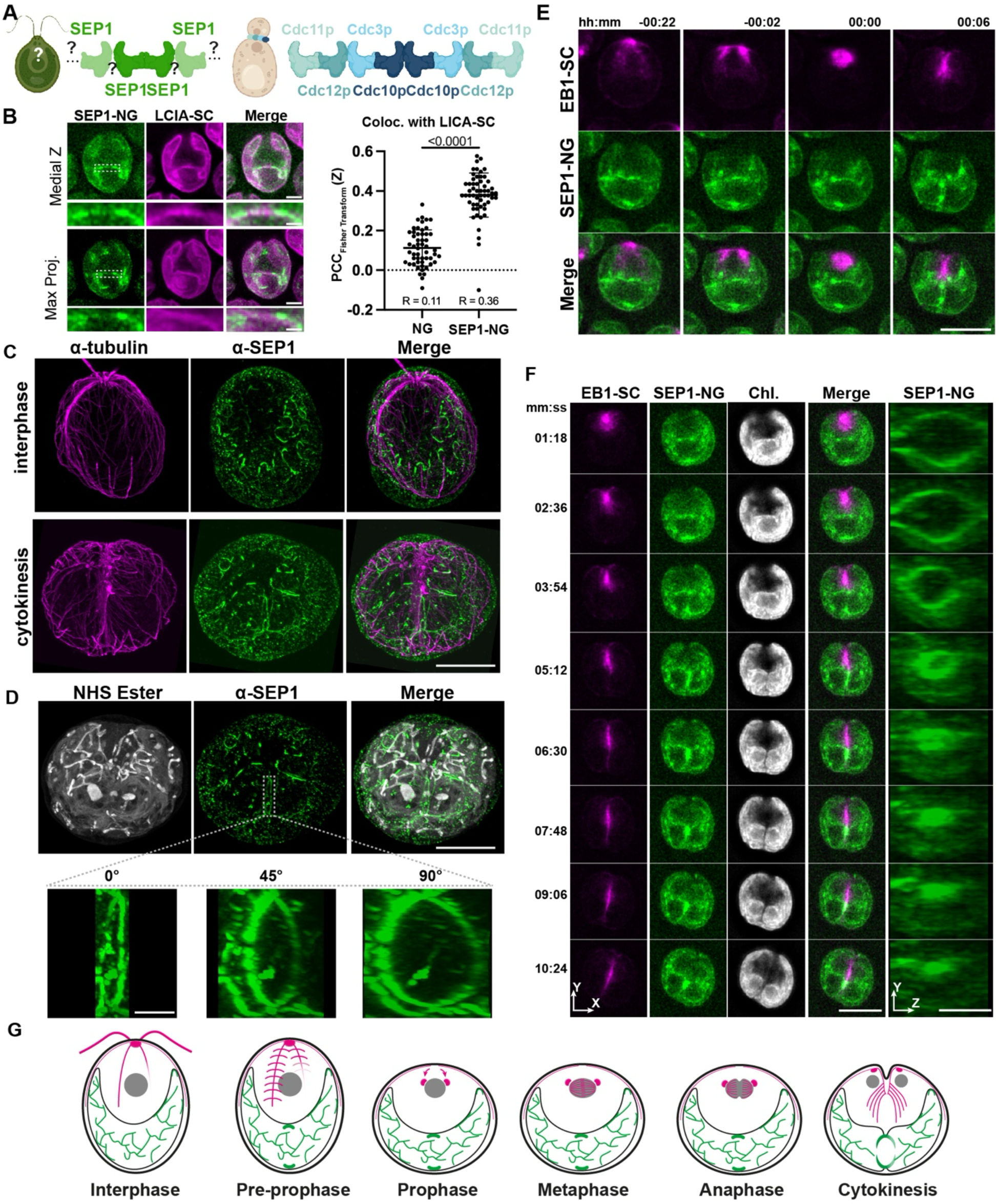
SEP1 localizes to the chloroplast surface and reorganizes into a division ring during cytokinesis. (A) Cartoon schematic of potential modes of septin dimerization/polymerization in *Chlamydomonas* compared to better-studied septins in yeast. (B) *Left,* SEP1-NG-3FLAG colocalizes with the chloroplast outer envelope marker LCIA-SC, forming small enrichments along the chloroplast surface in interphase. Representative medial Z-plane and max Z projection are shown. Scale bar, 3 μm. Insets, magnified region of chloroplast envelope. Scale bar, 1 μm. *Right,* quantification of NG/LCIA-SC and SEP1-NG/LCIA-SC colocalization using Fisher Z-transformed Pearson’s correlation coefficients (PCC). N = 53 (NG) and 56 (SEP1-NG). Unpaired Welch’s t-test. R indicates the back-transformed average PCC. (C) Ultrastructural expansion microscopy (U-ExM) images of anti-tubulin and anti-SEP1 staining in interphase and dividing cells showing filamentous networks of SEP1. Scale bar, 5 μm (adjusted for expansion) (D) *Top*, bulk protein labeling (NHS ester) localizes a SEP1 ring at the chloroplast division site. Scale bar, 5 μm. Bottom, 3D reconstruction of SEP1 signal rotated by the indicated angle along the Y-axis. Scale bar, 1 μm (adjusted for expansion). (E) Time-lapse imaging of SEP1-NG with the microtubule plus-end marker EB1-SC during mitosis. Scale bar, 10 μm. (F) Time-lapse imaging during cytokinesis showing constriction of the SEP1-NG ring at the chloroplast division site. Scale bar, 10 μm. The right-most panels show a magnified, 3D view of the SEP1 ring in the YZ-plane. Scale bar, 4 μm. (G) Interpretive diagrams of SEP1 (green) and EB1 (magenta) localization throughout the cell cycle.

Septins in opisthokonts form heteropolymeric filaments on membranes (See Introduction). Because of the evolutionary distance from opisthokonts and the presence of only one septin in *Chlamydomonas*, it was unclear whether the signal on the chloroplast outer envelope represented filaments. To address this, we employed ultrastructure expansion microscopy (U-ExM), in which cells were physically expanded by a factor of 4 to achieve an optical resolution equivalent to 44 nm. Using an antibody raised against *Chlamydomonas* SEP1, we found that the enrichments observed on the chloroplast surface during interphase are, in fact, filamentous networks (Fig. 1C, top; Supplementary Movie 1). A control experiment using a *sep1Δ* CRISPR mutant (see below) confirmed that these filamentous structures do not arise from non-specific staining of the antibody (Supplementary Fig. 2).

In dividing cells, portions of these networks reorganize into an apparent enrichment in the cell-division plane marked by microtubules (Fig. 1C, bottom). When compared with bulk-protein staining with NHS ester (which highlights the chloroplast among other structures), the position of the apparent enrichment coincided with the chloroplast-division site (Fig. 1D, top); 3D rendering and rotation of the images revealed that SEP1 filaments form a ring at this site (Fig. 1D, bottom; Supplementary Fig. 2; Supplementary Movies 2-3). Time-lapse imaging showed that the SEP1 ring formed during prophase and persisted at the cleavage plane through cytokinesis (Fig. 1E, Supplementary Movie 4), constricting as the cell and its single chloroplast divided (Fig. 1F, Supplementary Movie 5). This process of ring formation and constriction was repeated during every iteration of chloroplast division throughout the successive division cycles in *Chlamydomonas*^51^ (Supplementary Fig. 3; Supplementary Movie 6). Taken together, these findings provide the first *in vivo* evidence that the single septin in *Chlamydomonas* forms bona fide filaments and show that SEP1 dynamically reorganizes at the chloroplast envelope to form a medial ring involved in division (Fig. 1G).

### The amphipathic helix is required for SEP1 association with curved surfaces of the chloroplast envelope

The chloroplast surface in *Chlamydomonas* is geometrically heterogeneous, containing areas of both positive and negative principal curvature that dynamically reorganize during division ^52^. Septin filaments in animals and fungi associate with membrane surfaces of micron-scale curvatures in part through the amphipathic helix present in some subunits^50,53^. SEP1 contains a predicted C-terminal amphipathic helix (AH), a domain often associated with sensing micron-scale membrane curvature ^50,54^. Expression of full-length SEP1-NG-3FLAG in the *sep1Δ* background restored chloroplast localization patterns observed in the endogenously tagged strain (Supplementary Fig. 4A). Deletion of the AH domain (SEP1^ΔAH^-NG-3FLAG) resulted in context-dependent localization. When expressed in the WT background, SEP1^ΔAH^-NG-3FLAG localized to small puncta at the chloroplast periphery, with faint additional signal at the plasma membrane. However, when expressed in the *sep1Δ* background, chloroplast localization was abolished, and the fusion protein was fully directed to the plasma membrane (Supplementary Fig. 4A). Expression of the AH domain alone (AH_SEP1_-NG-3FLAG) was insufficient for chloroplast targeting (Supplementary Fig. 4B). These results indicate that the AH domain is necessary but not sufficient for chloroplast membrane association of SEP1 and suggest that additional domains or structural interactions are required. The partial recruitment of SEP1^ΔAH^-NG-3FLAG to the chloroplast envelope in the presence of endogenous SEP1 provides *in vivo* evidence that SEP1 self-associates, most likely through dimerization and polymerization, consistent with its ability to form higher-order structures as observed by U-ExM.

Septin filaments in budding yeast sense micron-scale membrane curvatures in a composition-dependent manner: increasing the number of AH domains per unit filament length enhances binding to highly curved surfaces ^50^. Shs1-capped septin filaments (Cdc3-Cdc10-Cdc12^AH^-Shs1^AH^), which contain an extra AH compared to Cdc11-capped filaments (Cdc3-Cdc10-Cdc12^AH^-Cdc11), preferentially associate with more strongly curved membranes ^54^. Although not yet systematically tested, a similar principle applies to mammalian septin complexes with varying numbers of AH domains ^35^. If we assume that *Chlamydomonas* SEP1 forms monomeric polymers, the density of AH domains is 2-4x higher than in yeast septin filaments. To quantify the structural properties of SEP1 filaments, we measured the lengths and curvatures of individual filaments. SEP1 filaments had an average length of 1.7 μm (σ = 0.73 μm, n = 24), suggesting that they can detect micron-scale curvatures. Curvature measurements using Menger’s method across varying window sizes revealed average Κ values (inversely proportional to the actual curvature) ranging from 2-6 μm^-1^ (Supplementary Fig. 4C–E), which is greater than a Cdc3-Cdc10-Cdc12^AH^-Cdc11 filament (∼2 μm^-1^), and more similar to Cdc3-Cdc10-Cdc12^AH^-Shs1^AH^ filaments (2-6.7 μm^-1^), consistent with the above idea.

### SEP1 is required for proper positioning of chloroplast division machinery

Given that SEP1 assembles into a medial ring at the chloroplast division site (Fig. 1), we hypothesized that SEP1 may play a functional role in chloroplast division. To test this, we used CRISPR/Cas-mediated genome editing to insert a hygromycin-resistance marker into a region of the *SEP1* locus encoding the central GTPase domain (Supplementary Fig. 5A-B). Genetic linkage analysis confirms that hygromycin resistance co-segregated with the *sep1Δ* allele (Supplementary Fig. 5C-D), and Western analysis using an anti-SEP1 antibody confirmed the loss of a ∼55kD band in the mutant line (Supplementary Fig. 5E). Our initial characterization of the *sep1Δ* mutant revealed no difference from WT in their growth or chloroplast morphology under standard growth conditions (Supplementary Figure 6). Thus, we speculated that septin plays an auxiliary role in chloroplast division, which may be revealed when the process is perturbed through an independent mechanism.

Previous work in *Chlamydomonas* has shown that, while F-actin is dispensable for cytokinetic furrow ingression, it is required for proper chloroplast division ^55,56^. Thus, we reasoned that F-actin perturbation may reveal the hidden role of septins in chloroplast division. Disruption of the actin cytoskeleton by Latrunculin B (LatB) in *Chlamydomonas* requires a *nap1* mutant background, in which the unconventional, LatB-resistant actin NAP1 is absent ^57–59^. We generated *sep1Δ nap1-1* double mutants and performed growth assays in the presence of a sub-lethal dose of LatB (0.5 μM) (Fig. 2A). While WT and *sep1Δ* strains showed no sensitivity, *nap1-1* showed moderate growth defects under LatB treatment, as expected. In contrast, the *sep1Δ nap1-1* double mutant exhibited a stronger synthetic growth defect; this sensitivity was rescued by transgenic expression of SEP1-NG-3FLAG (Fig. 2A), confirming that the phenotype was SEP1-dependent.

**Figure 2.**
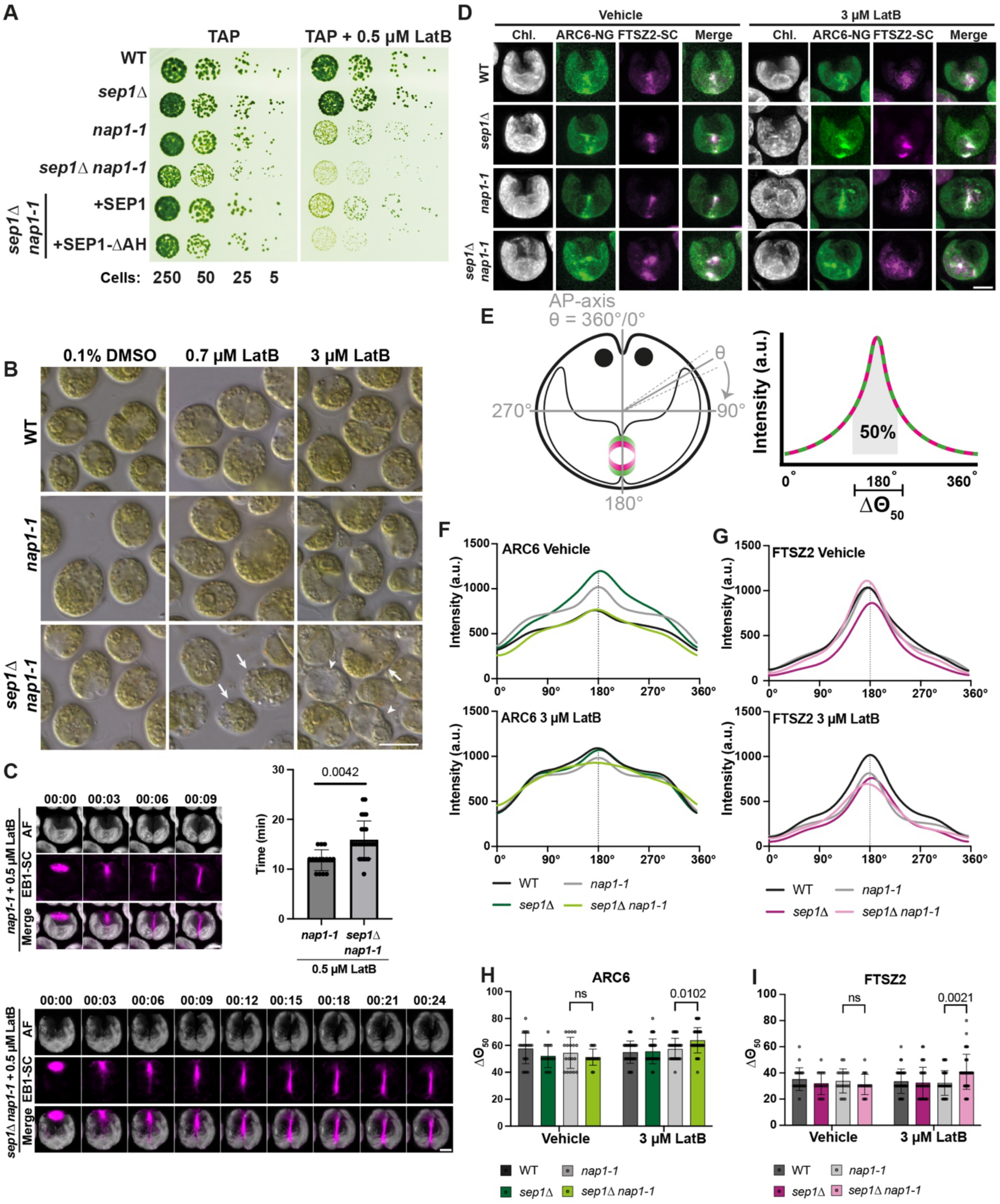
SEP1 is required for proper position of chloroplast division machinery. (A) Spot assay comparing growth of WT, *sep1Δ*, *nap1-1*, *sep1Δ nap1-1*, and *sep1Δ nap1-1* transformed with *SEP1-NG-3FLAG* or *SEP1^ΔAH^-NG-3FLAG,* with or without a sub-lethal dose of Latrunculin B (LatB; 0.5 μM). (B) Differential interference contrast (DIC) images of WT, *nap1-1*, and *sep1Δ nap1-1* cells treated with sub-lethal (0.7 μM) or lethal (3 μM) concentrations of LatB for 18 hours. Arrows point to cells with abnormal chloroplast morphology and vacuolization. Arrowheads point to cells lacking pigmentation. Scale bar, 10 μm. (C) Time-lapse series of dividing *nap1-1* and *sep1Δnap1-1* cells. The cells were treated with a sub-lethal (0.5 μM) dose of LatB for 10 min prior to the beginning of imaging. Furrow ingression was monitored with EB1-SC, and chloroplast abscission was monitored with chlorophyll autofluorescence (Chl.). Scale bar, 5 μm. *Top right,* quantification of time from spindle assembly to observed chloroplast abscission in *nap1-1* and *sep1Δ nap1-1* cells. n = 15 (*nap1-1*) and 26 (*sep1Δ nap1-1*). (D) ARC6-NG-3FLAG and FTSZ2-SC-PA localization in WT, *sep1Δ*, *nap1-1*, and *sep1Δ nap1-1* cells treated with and without LatB. Scale bar, 5 μm. (E) Schematic showing the quantification method used in panels F–K. Cells are aligned by their anterior–posterior (AP) axis as a proxy for the cleavage plane. Signal intensity is integrated outward from 180° until 50% of the total area under the curve is captured, yielding Δθ_50_. (F) Polar plots of ARC6-NG-3FLAG intensity distribution in untreated and LatB-treated cells. (G) Polar plots of FTSZ2-SC-PA intensity in untreated and LatB-treated cells. (H-I) Mean Δθ_50_ values from individual traces in (F) and (G). Group comparisons were assessed using one-way ANOVA followed by individual Šidák pairwise comparison.

We next examined whether this genetic interaction reflected a defect in chloroplast division. Using differential interference contrast (DIC) microscopy, we analyzed the morphology of WT, *nap1-1*, and *sep1Δ nap1-1* cells following treatment with either sub-lethal (0.7 μM) or lethal (3 μM) concentrations of LatB. While WT and *nap1-1* cells maintained a generally symmetric chloroplast structure under low-dose treatment, *sep1Δ nap1-1* cells displayed lobed, distorted, or mispositioned chloroplasts (Fig. 2B, arrows), suggesting that SEP1 contributes to chloroplast morphology and symmetry during division. Under higher-dose LatB treatment (3 μM), we also observed a subpopulation of *sep1Δ nap1-1* cells that appeared unusually pale or translucent by color DIC (Fig. 2B, arrowheads), a phenotype not seen in WT or single mutants. This loss of visible pigmentation may suggest broader defects in chloroplast maintenance or function. While the underlying cause remains unclear, pale or albino phenotypes have been reported in land plant mutants with impaired chloroplast protein import ^11,12^.

While chloroplast morphology defects in *sep1Δ nap1-1* cells suggested potential defects in chloroplast division, these observations were indirect. We first asked if chloroplast division was delayed in the *sep1Δ nap1-1* mutant treated with a low dose of LatB. Using EB1-mScarlet-I (SC) as a cell cycle progression marker, we measured the time from mitotic spindle assembly to chloroplast separation (Fig. 2C). We observed a slight increase in the time from mitotic spindle to chloroplast separation, and a small proportion of cells where chloroplast fission appeared to be delayed (Fig. 2C). To directly test whether the LatB hypersensitivity observed in *sep1Δ nap1-1* cells reflects a defect in the organization or function of the chloroplast division machinery, we examined the localization of FTSZ1, FTSZ2, and ARC6. FTSZ1/2 are prokaryotic tubulin homologs that co-polymerize into a division ring in the chloroplast stroma, and ARC6 is a transmembrane anchor that tethers the FTSZ ring to the inner envelope. In untreated cells, all proteins localized to the chloroplast midzone of all strains tested (Fig. 2D; Supplementary Fig. 7A). Upon 3 µM LatB treatment, *sep1Δ nap1-1* cells exhibited mispositioned ARC6-NG-3FLAG, FTSZ2-SC-PA, and FTSZ1-SC-PA signals, often displaced from the chloroplast center (Fig. 2D, Supplementary Fig. 7A). To quantify this mispositioning, we developed an angular integration analysis: cells were oriented along their anterior–posterior (AP) axis, and fluorescence intensity was measured along a polar coordinate system to obtain signal intensity as a function of θ. We next integrated outward from 180° until 50% of the total integrated signal was captured, yielding a Δθ_50_ value for each cell (Fig. 2E).

In LatB-treated conditions, *sep1Δ nap1-1* mutants displayed significantly increased Δθ_50_ values for ARC6-NG-3FLAG, FTSZ2-SC-PA, and FTSZ1-SC-PA compared to *nap1-1* single mutants (Fig. 2F–I, Supplementary Fig. 7B-G), indicating a broader, less focused distribution of division protein signal. No significant differences were observed between genotypes in untreated conditions. Taken together, these results demonstrate that SEP1 is required for the proper spatial positioning of stromal chloroplast division proteins and that this role becomes more important under conditions of F-actin disruption.

### SEP1 interacts with outer envelope translocon GTPases in late G1 phase

To identify SEP1-interacting proteins specific to the chloroplast envelope, we performed immunoprecipitation followed by mass spectrometry (IP-MS) using *sep1Δ* strains expressing full-length SEP1-NG-3FLAG and the mislocalized SEP1^ΔAH^-NG-3FLAG mutant (Supplementary Fig. 3D). Because SEP1^ΔAH^-NG-3FLAG is excluded from the chloroplast envelope, comparison of these two datasets enables spatial resolution of chloroplast-specific SEP1-associated proteins. Among the envelope-enriched interactors, we identified multiple components of the TIC/TOC (translocons at the inner/outer chloroplast membrane) and the Orf2971-FtsHi ultracomplex (Fig. 3A–B, Table S1-2). The TIC/TOC complex is a multi-subunit, megadalton-scale protein-import machinery that spans the inner and outer membranes of the chloroplast ^60,61^. Out of 21 TIC/TOC components in *Chlamydomonas,* 11 were found among the 19 proteins detected as significantly enriched (adjPval < 0.05) in the SEP1-NG-3FLAG immunoprecipitates (Fisher’s Exact Test P value < 0.0001 Fig. 3A, Table S2-3). Similarly, out of 20 Orf2971-FtsHi components ^62^, 4 were round enriched (P = 0.0001; Fig. 3A, Table S2-3).

**Figure 3.**
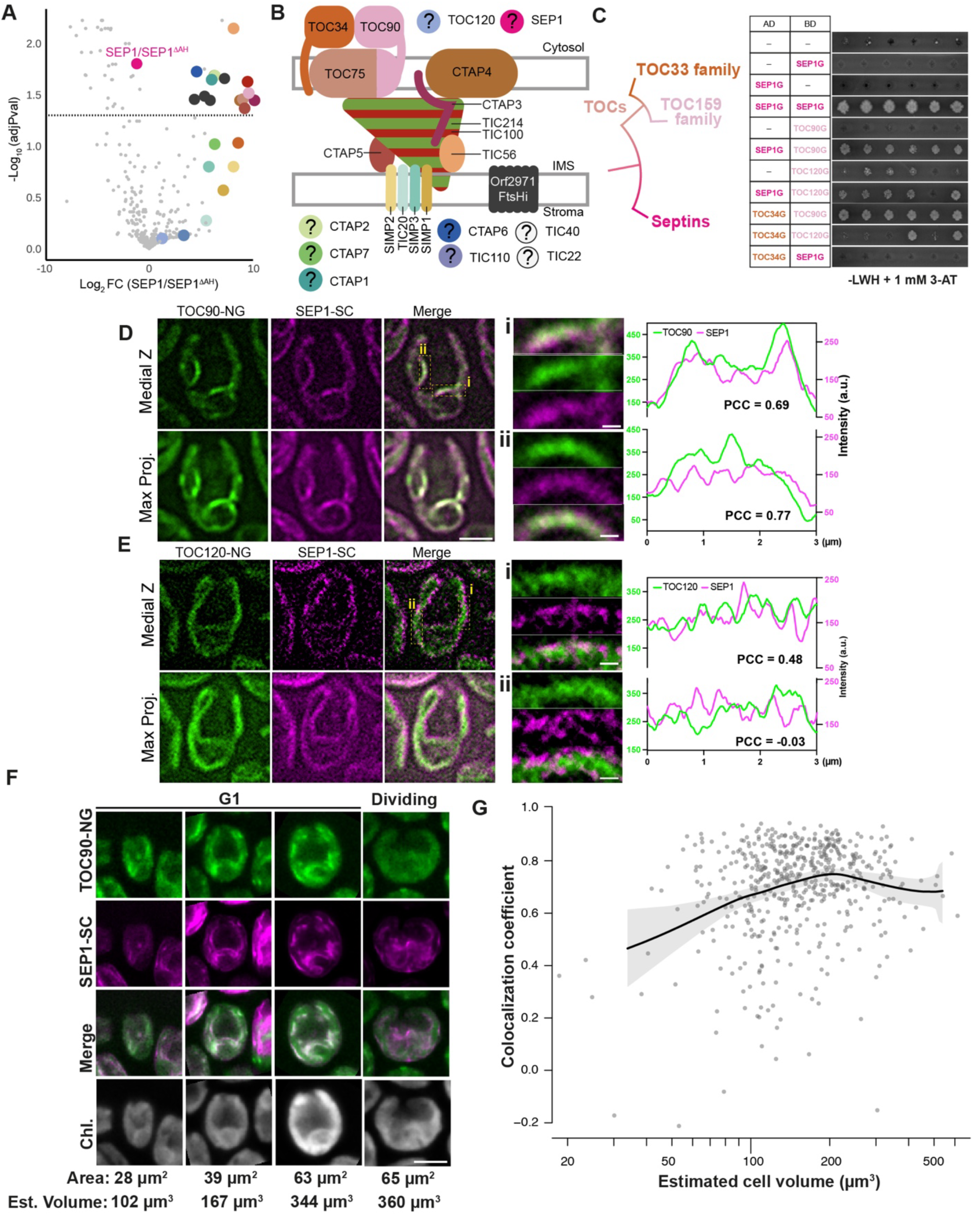
SEP1 binds to chloroplast translocons via a direct interaction with evolutionary related TOC GTPases. (A) Volcano plot of proteins identified by immunoprecipitation mass spectrometry from full-length SEP1-NG-3FLAG and SEP1^ΔAH^-NG-3FLAG (plasma membrane). Dotted line marks FDR-adjusted P = 0.05 (LIMMA empirical Bayes-moderated t-test with Benjamini-Hochberg correction). Colored points highlight components of the TOC-TIC complex and the Orf2971-FtsHi motor complex. Question marks indicate proteins identified as components of the TOC-TIC complex whose precise positions and roles remain unclear. IMS, inter-membrane space. (B) Schematic of the TOC-TIC complex and the Orf2971-FtsHi motor complex in *Chlamydomonas*. Colors are matched to those in (A). (C) *Left*, previously reported phylogenetic relationships among the GTPase domains of septins and TOC GTPases (Weirich et al., 2008). *Right*, yeast two-hybrid assay using the GTPase domains of SEP1, TOC34, TOC90, and TOC120. PJ69-4A cells were co-transformed with plasmids encoding SEP1-G fused to a Gal4 activation domain (AD) and TOC34-G, TOC90-G, TOC120-G, or SEP1-G fused to a DNA-binding domain (BD). Individual colonies represent independent co-transformants. Interaction was assessed on SC –Leu/–Trp/–His plates supplemented with 1 mM 3-AT. (D) Super-resolution images of a cell co-expressing SEP1-SC-3HA and TOC90-NG-3FLAG. Scale bar, 5 μm. (i-ii) Insets of individual SEP1-TOC90 enrichments. Scale bar, 0.5 μm. Line traces of SEP1 and TOC90 intensities and PCC are shown at *bottom*. Scale bar, 1 μm. (E) Super-resolution images of a cell co-expressing SEP1-SC-3HA and TOC120-NG-3FLAG. Scale bar, 5 μm. (i-ii) Insets of individual SEP1-TOC90 enrichments. Scale bar, 0.5 μm. Line traces of SEP1 and TOC120 intensities and PCC are shown at *bottom*. Scale bar, 1 μm. (F) Representative images of cells co-expressing SEP1-SC and TOC90-NG-3FLAG during G1 growth and dividing cells. Cell volumes are estimated by assuming an ellipsoidal geometry with an aspect ratio of 1.2 between length and width. Scale bar: 5 μm. (G) PCC values from individual cells plotted as a function of estimated cell volume. Means and 95% CIs of 1,000×-bootstrapped locally estimated scatterplot smoothing (LOESS) curves are shown.

These complexes are together responsible for the import of most nucleus-encoded chloroplast proteins: Import initiation is mediated by two classes of transmembrane GTPases: TOC33-family and TOC159-family receptors. In *Chlamydomonas*, these receptor families consist of a single TOC33-family member (TOC34) and two TOC159-family members (TOC90 and TOC120).

These proteins heterodimerize and recognize the N-terminal chloroplast-targeting peptides (cTPs) of cytosolic preproteins and initiate their translocation into the chloroplast stroma, where the Orf2971-FtsHi ATPase motor pulls the preproteins, and then stromal processing peptidases cleave cTPs ^63–65^. The interaction between SEP1 and the TIC-TOC complex was also confirmed by an orthogonal proximity-labeling approach using BioID (see below).

Given the enrichment of TOC34 and TOC90 in SEP1 immunoprecipitates, we tested whether SEP1 directly interacts with these receptors via their respective GTPase domains. Previous structural studies have shown that TOC GTPases share distant homology with septins ^66^, raising the possibility that SEP1–TOC interactions occur through conserved G-domain interfaces (Fig. 3C). Using yeast two-hybrid (Y2H) assays, we tested pairwise interactions between the GTPase domain of SEP1 (SEP1G) and those of TOC34, TOC90, and TOC120.

SEP1G interacted robustly with TOC90G and TOC120G, but not with TOC34G (Fig. 3C), suggesting specificity for TOC159-family receptors. In addition, we observed strong growth in SEP1G–SEP1G pairs, supporting our previous conclusion (Supplementary Fig. 3) and prior evidence that SEP1 is capable of homodimerization ^45^.

In canonical models, septins polymerize via two interfaces, the G- and NC-interfaces, giving rise to their characteristic palindromic structure. To better understand how a SEP1-TOC90 interaction might look, we utilized a published AlphaFold-based tiling method called FragFold. Using a moving window of 30 amino acid fragments from SEP1, we used FragFold to calculate individual weighted contact scores for both a SEP1-SEP1 homodimer and a SEP1-TOC90 heterodimer (Supplementary Fig. 8A-B). Many regions important for SEP1-SEP1 homodimerization were mapped on β-meander, a-4 helix, and the R-finger loop within the G-interface, as well as the a2-β4 loop within the NC-interface, consistent with SEP1’s ability to form homopolymers (Supplementary Fig. 8C). Similarly, many of the putative SEP1-TOC90 contact sites belong in the G-interface (Supplementary Fig. 8D); unlike with SEP1-SEP1, the a2-ý4 loop did not score high as SEP1-TOC90 contact site, potentially suggesting that the interaction between these two proteins are limited to the G-interface. We then asked whether SEP1G and TOC34G could compete against each other for binding to TOC90G. Using the same FragFold approach, we tiled the interaction scores for both SEP1G and TOC34G across TOC90G. While the predicted binding profiles were clearly distinct, some regions overlapped in the GTP-binding face of TOC90G, suggesting a potential competition (Supplementary Fig. 8E-H). Further biochemical examinations are required to understand the exact nature of this interaction.

Next, we tested the colocalization between SEP1 and the TOC GTPases in vivo. TOC90-NG showed discrete enrichment along the chloroplast envelope that largely overlapped with SEP1-SC, with overall high colocalization coefficients (Fig. 3D). The TOC120-NG signal was noticeably weaker than that of TOC90-NG (both expressed using their native promoters), and colocalization with SEP1-SC was less strong (Fig. 3E). Because previously published proteomic experiments on the *Chlamydomonas* translocon did not detect TOC120 ^67^, TOC90 may be the major TOC159-family GTPase under the conditions used. Using the TOC90-NG SEP1-SC pair, we asked whether the timing of colocalization is controlled relative to the cell-division cycle.

While the SEP1-SC signals remained strongly associated with the chloroplast envelope forming filaments and a ring at the division site, the signal of TOC90-NG was detectable only in G1 cells (Fig. 3F). When the colocalization coefficient between the two proteins was plotted as a function of estimated cell volume (assuming a spheroidal cell shape with the aspect ratio of 1:1.2), it gradually increased until a peak at ∼200 µm^3^, after which it began to decline (Fig. 3G). This peak coincides with the cell volume just before cell division, according to the diurnal transcriptome study by Zones et al ^68^; see also: Fig. 4B). These results suggest that TOC90 and SEP1 interact with each other on the chloroplast envelope throughout the long G1 phase in *Chlamydomonas*. In dividing cells, while TOC90 is removed from the chloroplast through an unknown mechanism, SEP1 remains associated to form a ring. Whether the SEP1 ring contains other TOC components or it functions independently of the translocon complex remains unclear.

**Figure 4.**
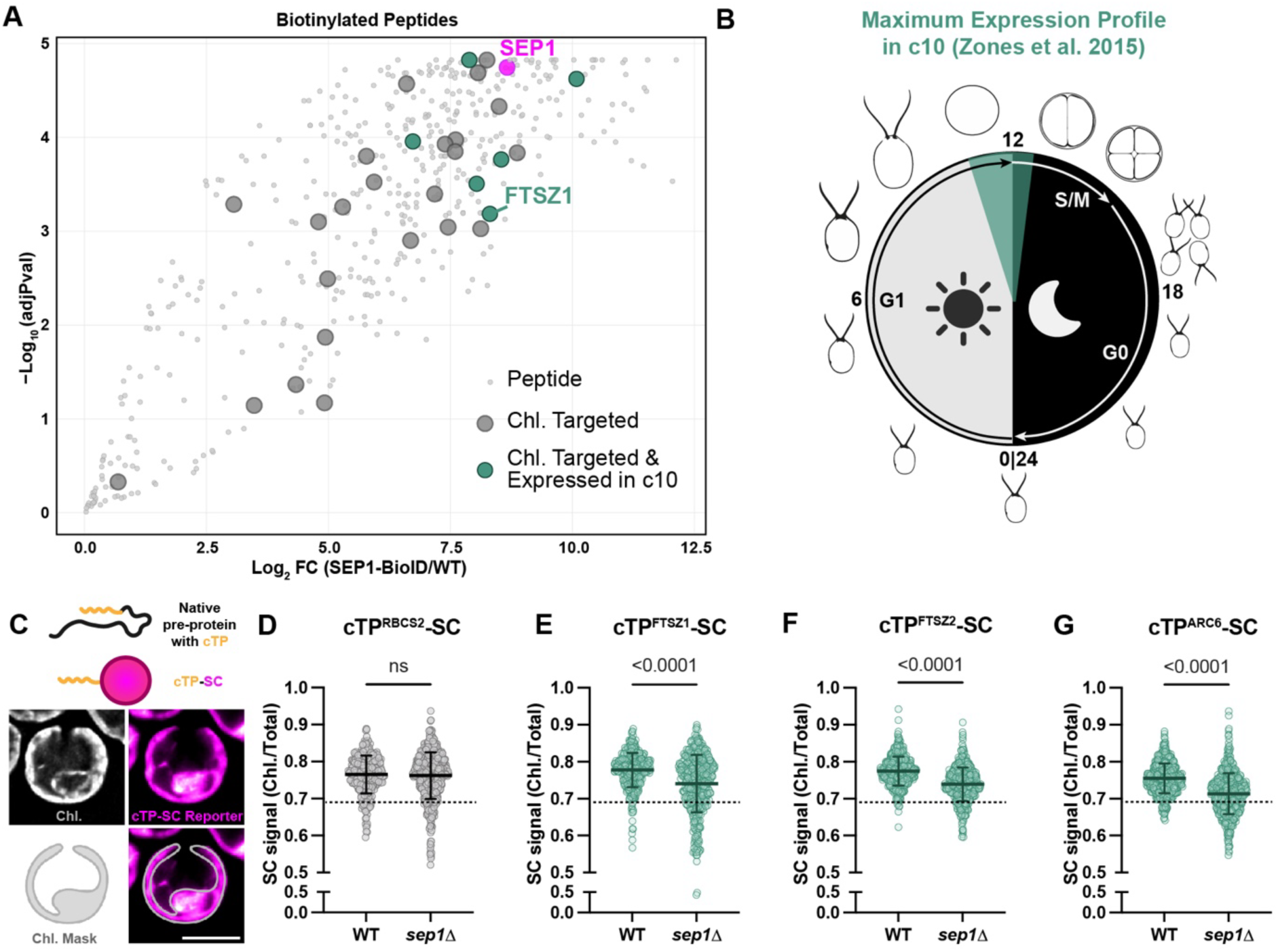
SEP1 enhances import of chloroplast division proteins. (A) Volcano plot of peptides biotinylated by SEP1-NG-BioID. Larger points represent proteins computationally predicted to be chloroplast-targeted. Teal points represent peptides from proteins whose maximal gene expression occurs during division. (B) *Chlamydomonas* diurnal cell cycle. Teal region corresponds to the window for maximal expression surrounding the division cycle based on Zones et al. 2015. (C) Schematic diagram of translocation reporters. Chloroplast targeting peptides (cTP) of specific proteins were fused to SC fluorescent protein. The fraction of signal in the chloroplast over the total signal in the cell was measured using a chloroplast mask generated from chlorophyll autofluorescence. Scale bar, 5 μm. (D-G) Fractions of chloroplast signal using reporters containing cTPs from (D) RBCS2, (E) FTSZ1, (F) FTSZ2, and (G) ARC6. Dotted lines represent baseline fluorescence determined from the control cycloheximide experiment shown in Supplementary Figure 10.

### SEP1 interacts with chloroplast-targeted proteins

To gain further insight into the role of SEP1 on the chloroplast envelope, we performed a BioID-based proximity-labeling experiment using a transgenic SEP1-NG-BioID construct. This construct localized to the chloroplast surface and was enriched at the chloroplast division site (Supplementary Fig. 9A). Synchronous cultures of cells were treated with 1 mM biotin for 24 hours, and biotinylated proteins were pulled down using streptavidin beads. This experiment identified both TOC90 and TOC120, along with other known translocon components, consistent with the previous IP-MS and Y2H data (Supplementary Fig. 9B-C, Table S4).

By filtering individual biotinylated peptides from proteins that were computationally predicted to be chloroplast targeted, we identified the stromal division protein FTSZ1 as a top biotinylated candidate, among other chloroplast-targeted proteins whose gene expression is maximal around the onset of division (Fig. 4A-B). Because we did not identify FTSZ1 or any other chloroplast-division proteins in the SEP1 co-immunoprecipitation mass-spectrometry experiment, we speculated that this biotinylation signal may reflect a transient interaction between SEP1 and FTSZ1 preprotein as the latter is translocated into the chloroplast. To test this idea, we expressed reporter proteins consisting of the N-terminal chloroplast targeting peptides (cTPs) from RBCS2 and FTSZ1 fused to mScarlet (SC) in WT and *sep1Δ* backgrounds (Fig. 4C). RBCS2 is a Rubisco small subunit protein constitutively expressed throughout the cell cycle ^68^, and the cTP^RBCS2^-SC reporter localized in the chloroplast in both WT and *sep1Δ* cells (Fig. 4D). When the cTP^RBCS2^-SC cells were treated with a potent translation inhibitor, cycloheximide, the chloroplast/total cell cTP^RBCS2^-SC signal ratio plateaued at the ∼0.69 bottom (Supplementary Fig. 10). This high remaining ratio despite the complete blockage of protein synthesis is likely due to the strong background autofluorescence of the chloroplast that occupies >50% of cell volume ^52^. When cTP^FTSZ1^-SC was examined, there was a statistically significant decrease in the ratio from WT cells compared to *sep1Δ* cells (Fig. 4E), equating to nearly 50% of the effects of cycloheximide. Similar effects were observed for two other preproteins of chloroplast-division proteins examined, FTSZ2 and ARC6 (Fig. 4 F-G), suggesting the general importance of SEP1 in translocation of chloroplast-division proteins.

To explore potential mechanisms underlying the observed import defect, we considered several alternative hypotheses. One possibility was that SEP1 is required for spatial patterning of TOC90 relative to the chloroplast division site. Imaging of dividing cells does not show a noticeable effect of *sep1Δ* on the TOC90 signal, which is absent from the chloroplast division site (Supplementary Fig. 11A). Another hypothesis is that SEP1 may be important for global patterning of translocon components during interphase. Imaging WT and *sep1Δ* cells expressing either TIC20-Venus or TOC90-NG-3FLAG did not show changes in the patchy distribution of signal on the chloroplast membrane, arguing against large-scale mispositioning of chloroplast translocons as the cause of impaired import (Supplementary Fig. 11B-C). Together, these results indicate that SEP1 does not regulate TOC localization. Instead, they are consistent with a model in which SEP1 promotes the efficient import of specific substrates, either by acting as an effector to regulate translocon substrate selectivity or by acting directly as a novel subunit of the chloroplast translocon itself.

### TOC GTPases may have evolved from an ancient septin

Our data so far suggest a model in which SEP1 interacts with TOC GTPases via conserved G-interfaces and promotes efficient import of division proteins. The previous molecular phylogenetic analysis showing similarity but a clear bifurcation between septins and TOC GTPases included only septins from opisthokonts and TOC GTPases from land plants ^69^. Thus, we revisited their evolutionary relationship using the expanded availability of septin sequences across the eukaryotic tree. We performed a maximum-likelihood phylogenetic analysis using representative septins from opisthokonts, algae, and protists, along with TOC GTPases across Archaeplastida, including green algae and early-diverging red algae (Table S5). The resulting topology revealed that TOC-family proteins form a distinct monophyletic clade that branches within the broader septin family (Fig. 5A, Supplementary Fig. 12). Notably, TOC sequences clustered most closely with septins from early-branching algae, supporting the view that TOC GTPases derived from an ancient algal septin subfamily. Sequence alignment of SEP1 with TOC34 GTPases also demonstrate shared conservation of arginine finger (R239 in SEP1), a residue that is critical for their dimerization and GTPase activity (Fig. 5B). This new phylogenetic relationship places chloroplast translocons as a sub-clade of septins following the speciation events between opisthokonts and non-opisthokonts (Fig. 5C). The emergence of the chloroplast translocon was a critical cellular innovation for the Last Archaeplastida Common Ancestor to transform an ancient cyanobacterium into a modern chloroplast. Septins were likely once a molecular feature of this common ancestor, although they may have been lost in later diverging land plants (Fig. 5D).

**Figure 5.**
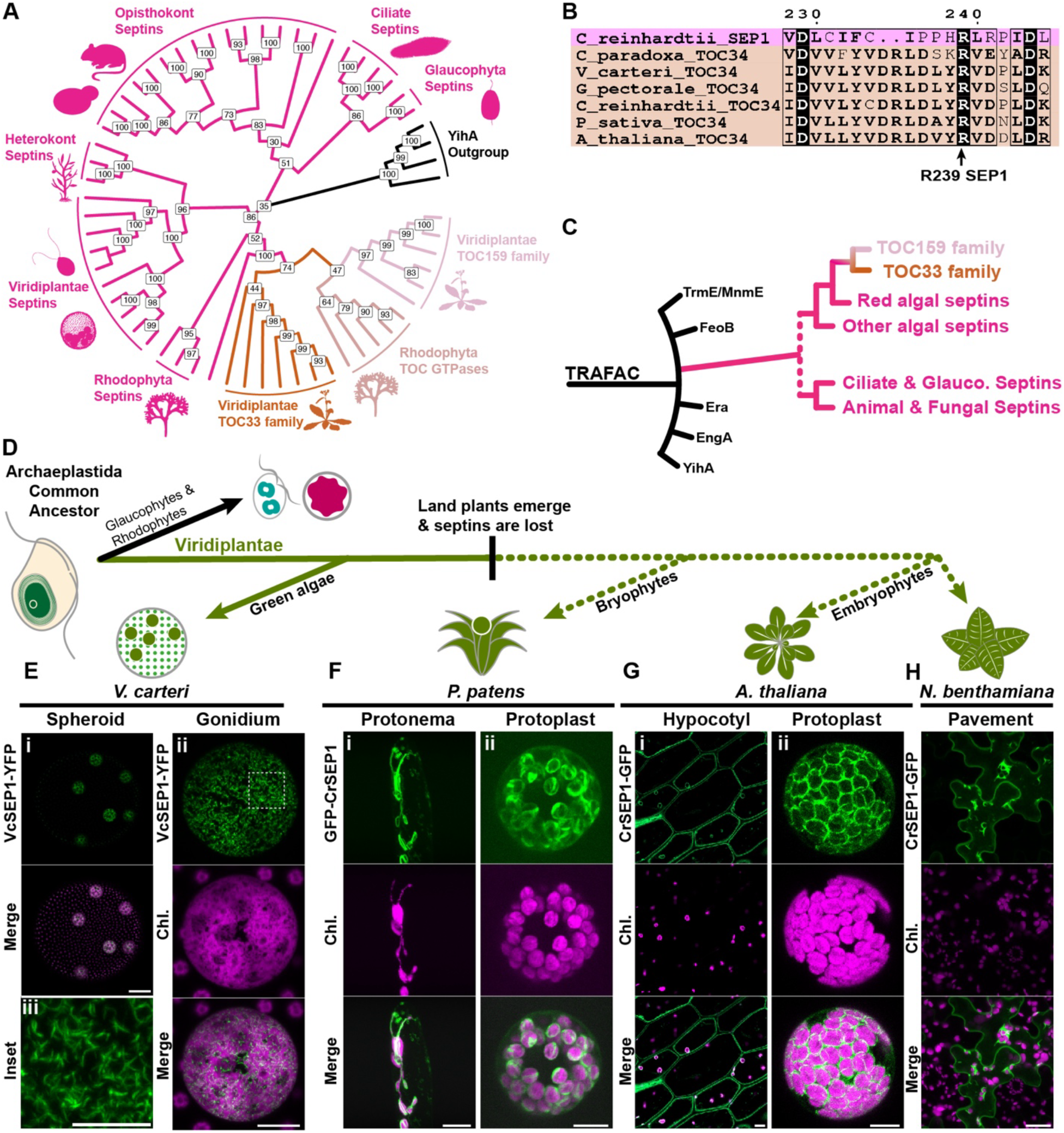
**TOC GTPases are evolutionarily derived from an ancestral algal septin**. (A) Cladogram of the best maximum likelihood tree (see Supplementary Figure 12) showing the relationship between septins and Archaeplastida TOC GTPase proteins. Bootstrap support values are shown at nodes. Bacterial YihA-family GTPases were used as the outgroup. (B) Multiple sequence alignment of SEP1 and TOC34 GTPase domains. Conserved arginine-finger residue is highlighted. (C) New proposed model of septin and TOC family GTPase evolution, based on Weirich et al., 2008 and (A). (D) Representative timeline of Archaeplastida evolution and septin gene loss. (E) Localization of YFP-tagged single *Volvox carteri* septin VcSEP1. (i) Chloroplast envelope enrichment in a single spheroid. Scale bar: 100 μm. (ii) Single gonidial chloroplast. Scale bar: 20 μm, (iii) Inset of gonidial chloroplast. Scale bar, 10 μm. (F-H) Heterologous expression of GFP-tagged *Chlamydomonas* CrSEP1 in various plant species. Scale bars, 10 µm. (F) *Physcomitrium patens*. (i) Protonema. (ii) Protoplast. (G) *Arabidopsis thaliana.* (i) Hypocotyl cells. (ii) Protoplast. (H) *Nicotiana benthamiana* pavement cells.

To determine whether SEP1 retains conserved chloroplast targeting capacity across evolutionary distance, we turned to *Volvox carteri*, a multicellular relative of *Chlamydomonas*. We expressed the single *V. carteri* septin, VcSEP1, as a fusion with YFP. *V. carteri* consists of two cell types: large, actively dividing gonidial cells surrounded by a monolayer of terminally differentiated somatic cells. VcSEP1-YFP localized to the chloroplast surface in the actively dividing gonidial cells, but barely detectable in non-dividing somatic cells. (Fig. 5E).

Next, to test whether the chloroplasts of land plants have retained the ability to be recognized by septin, despite the universal loss of this gene family. We examined three species spanning major transitions in plant evolution: the early-diverging moss *Physcomitrium patens*, and the later-diverging angiosperms *A. thaliana* and *Nicotiana benthamiana*. In all systems, heterologously expressed CrSEP1 localized to the chloroplast periphery in both whole tissue and isolated protoplasts (Fig. 5F-H). In *P. patens*, GFP-CrSEP1 formed lasso-like filaments and complete rings along the chloroplast surface (Fig. 5Fi), and arc-like filaments coinciding with membrane indentations in isolated protoplasts (Fig. 5Fii). In *A. thaliana*, CrSEP1-GFP was also targeted to the chloroplast surface (Fig. 5Gi-ii). In *N. benthamiana*, CrSEP1 formed needle-like filaments preferentially associated with the chloroplast envelope, though occasional plasma membrane localization was also observed (Fig. 5H). These observations indicate that SEP1 retains chloroplast membrane-targeting capacity despite more than 700 million years of Viridiplantae evolution.

Finally, to test whether CrSEP1 retains its interaction with TOC components in heterologous systems, we performed immunoprecipitation followed by mass spectrometry in *N. benthamiana* leaf tissue expressing CrSEP1-GFP or GFP alone. Several TOC translocon GTPases were enriched in the SEP1 pulldowns, including members of the TOC159-family (Supplementary Fig. 13A-B, Table S6-S7), consistent with conserved recognition of TOC proteins in land plant chloroplasts. In addition to the TOC GTPases, our pulldowns also identified cTP-containing proteins. Many of these chloroplast-targeted proteins are division-related (Supplementary Fig. 13C-D). This was surprising because such interactions should normally be transient, and we indeed did not detect cTP-containing proteins by IP-MS in *Chlamydomonas.* It is possible that heterologous overexpression of CrSEP1 in *N. benthamiana* impairs chloroplast import, thereby enhancing detection of chloroplast-targeted proteins. When focusing specifically on chloroplast division proteins, we detected only cTP-containing components (FTSZ, ARC6, MIN), but not cytosolic proteins (DRP5B or PDV) (Supplementary Fig. 13D). This model further supports septin’s role in regulating chloroplast division via chloroplast translocation.

Together, these results reveal that TOC GTPases are phylogenetically derived from an ancestral algal septin, and that SEP1 retains both membrane-targeting and protein-interaction properties across a broad range of photosynthetic eukaryotes. This evolutionary continuity suggests that the functional relationship between septins and the chloroplast protein import machinery was established early in the evolution of Archaeplastida and has been retained across major transitions in Viridiplantae, in concert with lineage-specific adaptations.

## DISCUSSION

### Septin filaments on the chloroplast envelope

Although the presence of species outside of opisthokonts with single septins was pointed out by Verselle and Thorner as early as 2005 ^49^, the architecture and function of non-opisthokont septins remained understudied. By combining live-cell imaging with U-ExM, we found that the single *Chlamydomonas* septin forms bona fide filaments on the chloroplast surface (Fig. 1). This result is unexpected and significant in two ways: First, most current models of septin assembly are based on studies in the animal and fungal systems, in which multiple septin subunits assemble into heteromeric complexes and filaments. Second, association of septins with chloroplasts has not been reported, and together with the seemingly complete absence of septins at the cell-division site, this localization provides a new evolutionary perspective on the evolution of septins (see below).

Targeting of SEP1 to the chloroplast may involve multiple mechanisms. Our results show that targeting requires the C-terminal amphipathic helix (AH), although this domain alone is not sufficient. AH domains are key regulators of septin localization in other systems, potentially via a mechanism that recognizes specific membrane curvature^50^. Individual SEP1 filaments form curvatures more in line with Shs1-capped filaments from yeast, which contain more AH domains than Cdc11-capped filaments^54^. The chloroplast of *Chlamydomonas* is fenestrated and thus displays a wide range of potentially compatible membrane curvatures ^52,70^. Thus, curvature-sensing may be one mechanism by which SEP1 is targeted to the chloroplast. In addition, SEP1 may be targeted to the chloroplast by protein factors. Our data show that chloroplast-localized SEP1 binds almost exclusively to components of the TOC-TIC-Orf2971-FtsHi ultracomplex (Fig. 2) ^61,71^, and direct interaction was detected for TOC90. Whether this interaction is involved in the chloroplast targeting of SEP1 remains unclear. Attempts to generate *toc90Δ* deletions have been unsuccessful, likely indicating that this gene is essential for chloroplast import and cell viability.

The observations that SEP1 can form filaments *in vivo* (this study) and *in vitro* suggest that these filaments are homopolymeric ^43,45^. Indeed, the resolved crystal structures of SEP1 display conservation of motifs critical both for G- and NC-interface interactions. In humans, SEPT2 can form homotypic filaments with a single amino acid substitution *in vivo* ^72^. Similarly, mis-expressed individual septins can form homopolymers *in vivo* ^32,73,74^ such as human Sept9 whose isoforms are often overexpressed in tumors ^75^. Single septins in algae provide a unique system to investigate the molecular mechanism of homopolymer formation in a physiological, non-pathogenic context.

### Septin interacts with chloroplast translocon GTPases

Our results demonstrate that SEP1 interacts with chloroplast translocons at the chloroplast surface, with evidence supporting a direct interaction with TOC90. FragFold predictions place this interaction in the G-interface of SEP1, which is notable because TOC33-family and TOC159-family proteins also interact through their G-interfaces^63^. GTP hydrolysis by the TOC GTPases serves as an important checkpoint in preprotein sorting into the chloroplast ^76^. Biochemical studies of SEP1 have reported an unusually high GTPase activity when compared to other studied septins ^45^. Together, these observations raise the possibility that SEP1’s high GTPase activity is mechanistically linked to its role in chloroplast translocation.

The modest phenotype of the *sep1Δ* mutant may reflect functional overlap with other chloroplast GTPases. In land plants, distinct TOC33-family paralogs show both specialization and redundancy in their substrate preprotein recognition, and combined receptor mutants often produce much stronger defects than single mutant^12,26,77^. By analogy, SEP1 and TOC34 may provide partially overlapping activity in *Chlamydomonas*. This could explain why loss of SEP1 alone does not completely abolish chloroplast function. Assays for chloroplast protein import in the *sep1Δ* mutant also support this conclusion: While import of chloroplast-division proteins was affected, such an effect was not observed for an RBCS2, indicating that SEP1 possess some level of substrate selectivity. Our repeated attempts to generate a *toc34Δ* mutant using CRISPR were unsuccessful, suggesting that TOC34 may have broader substrate specificity, including some essential proteins.

These observations also suggest a structural model for how SEP1 may organize at the chloroplast surface. SEP1 could first engage TOC90 through its G-interface and then extend outward through alternating NC- and G-interface-mediated self-assembly, allowing TOC90 to act as a nucleator or anchor for filament growth. Alternatively, together with SEP1, the TOC complex itself may form filament-like distribution on the chloroplast surface. Live-cell, super-resolution, and/or ultrastructural imaging will be important for testing these models.

### The septin ring at the chloroplast division site

Septins in animals and fungi are known to form rings at sites of membrane curvature where they act as molecular scaffolds for protein recruitment and spatial organization. In *S. cerevisiae*, septins form a ring at the mother-daughter bud neck to recruit critical cytokinetic proteins to facilitate recruitment of cell-division proteins ^78–80^. In metazoa, septins are universally found at the cell-division site, such as in *Drosophila melanogaster*, *C. elegans,* and mammalian cells ^74,81–83^.

Our results extend the conservation of septin rings to outside of the opisthokont lineage. Time-lapse microscopy and U-ExM imaging confirmed that a filamentous SEP1 ring forms during mitosis. In two other green algae, *N. bacillaris* and *M. geminata*, their respective single septins were localized at the cell-division site by immunofluorescence studies ^43^. The chloroplasts in these small (1-2 µm), monoplastidic species are positioned medially during cell division, and it is unclear whether this localization occurs at the cytokinetic furrow or at the chloroplast division site. It should be interesting to revisit these reported septin localizations using higher-resolution imaging methods such as U-ExM.

Interestingly, SEP1 and TOC90 do not appear to colocalize at the division ring in mitotic cells. This spatial separation suggests that SEP1 may perform at least two distinct functions: one associated with TOC90 and the protein import machinery during late G1 phase, and another at the chloroplast division site during mitosis. The function of the mitotic SEP1 ring remains unclear, but one attractive possibility is that it serves as a scaffold for cytosolic chloroplast-division factors. In land plants, the dynamin-related protein DRP5B contributes to chloroplast division, and septins in mammalian cells can promote mitochondrial fission by organizing recruitment of dynamin-family proteins ^38,84^. The SEP1 ring may analogously help recruit or stabilize DRP5B-like machinery at the chloroplast division site. A second possibility is that the SEP1 ring helps coordinate chloroplast division with cytokinesis. *Chlamydomonas* cytokinesis depends on both microtubules and F-actin, and furrow-associated microtubule arrays approach the chloroplast division plane during mitosis ^85–87^. In this context, the SEP1 ring could act as a cytoskeletal landmark or crosslinking scaffold that helps align chloroplast constriction with ingressing cytokinetic structures. Because ring formation is cell-cycle restricted, identifying its upstream regulators will likely be essential for defining its function. Kinases or other mitotic factors that control SEP1 ring assembly may provide an entry point into this problem.

### The possible roles of ancient septins

Broad phylogenetic distribution of septins and the presence of species with single septins support an early origin in eukaryotic evolution as a single gene at or before LECA ^48^. A recurring feature of septins across eukaryotes is their assembly into higher-order structures, including filaments and rings, that are associated with membranes ^88^. These shared properties suggest that the ancestral septin may have been a membrane-associated GTPase capable of polymerizing into curved or ring-like assemblies. Our findings extend this framework by showing that a green algal septin associates with the chloroplast envelope, forms filaments, and reorganizes into a ring at the chloroplast division site. Considered alongside septin localization to mitochondrial membranes in ciliates and animals ^38,44,89^ and to intracellular bacteria in animal systems ^42^, our results suggest that binding to the surfaces of bacteria or bacterium-derived organelles and regulating their fission is a conserved septin function. One attractive possibility is that LECA septin was part of an ancient cell-autonomous defense system against intracellular bacteria, a eukaryote-specific problem.

Our findings also bear on the mysterious evolutionary origin of chloroplast TOC GTPases ^28^. The phylogenetic relationship between septins and TOC receptors, together with the direct interaction we observe between SEP1 and TOC90, supports a model in which these proteins retain both structural and functional continuity. One possible scenario is that, during early plastid establishment, an ancestral septin recognized the surface of the cyanobacterial endosymbiont and was subsequently adapted into a dedicated translocon-associated GTPase. In this model, acquisition of a transmembrane segment and stable association with prokaryotic surface proteins would have enabled this lineage to transition from a peripheral membrane organizer to a core component of the protein import machinery ^90^. This transition would have been essential for host control over the nascent organelle by allowing nuclear-encoded proteins, including chloroplast division factors, to be imported back into the endosymbiont-derived compartment.

## MATERIALS AND METHODS

### *C. reinhardtii* growth conditions and genetic crosses

Routine cell culture was done as previously described in Tis-acetate-phosphate (TAP) medium at ∼25°C under constant illumination at 40 μmol photons×m^-2^×s^-1^ ^55^. LatB was purchased from Adipogen (Cat. #: AG-CN2-0031, Lot #s: A01191/M and A01191/O) and dilutions into TAP medium were made from a 10 mM stock in dimethyl sulfoxide (DMSO). Genetic crosses were performed as described previously ^91^. When necessary, segregants were genotyped based on known phenotypes (selectable marker, fluorescence, LatB sensitivity) or by PCR using appropriate primers.

### Transformation

Transformation was always performed into the CC-124 WT strain and subsequently crossed into the noted genotype, so that transgenes were located at the same genomic locus in all comparisons. 1 μg of plasmid DNA was linearized using an appropriate restriction enzyme at a site outside of the expression cassette, in a 10 μl reaction (1 μl rCutSmart, 0.5 μl of restriction enzyme, volume to 10 μl with molecular grade H_2_O). Transformation was done by square-pulse electroporation using a NEPA21 electroporator ^92^. Transformants were selected for resistance to paromomycin (RPI, P11000; 5 μg/ml) or hygromycin B (VWR, 97064-454; 10 μg/ml) and subsequently for the expression of fluorescently tagged proteins.

### CRISPR-Cas9 mediated tagging and knockout

We designed guide RNA sequences using CHOPCHOP. The single guide RNAs (sgRNAs) were generated to target the sequence: 5’-CGACGAGCCGATGAGTGACAGGG (Cre12.g556250, *SEP1*). Donor DNA fragments were generated by PCR using primers SDP014 and SDP017 from plasmids pSD077 (for NG-3xFLAG tagging) or pSD078 (for a SC-PA tagging). These fragments contain sequences of the C-terminal end of SEP1 (from the CRISPR cut site to the last coding codon), a fluorescent marker and epitope tag, along with an *AphII* hygromycin-resistance cassette for selection. To generate RNP complexes, 120 pmol of IDT Alt-R CRISPR-Cas9 sgRNA was added to 61 pmol of Alt-R S.p. Cas9 Nuclease V3 (Integrated DNA Technologies). The final volume of this reaction was brought to 5 μl with IDT Duplex buffer and incubated at 37°C for 30 minutes.

CC-124 cells were grown in liquid TAP at 25°C under continuous light to ∼5 × 10^6^ cells/ml. Cells were collected and resuspended in 1 ml of autolysin media and incubated for 1 hour at room temperature. Cells were then centrifuged briefly (2,000 x *g,* 2 minutes) and resuspended at 2 × 10^8^ cells/ml in 0.5X TAP + 80 mM sucrose. For each reaction, the RNP together with 1 μg donor DNA was mixed with 125 μl of the autolysin-treated cell suspension and delivered by electroporation using a NEPA21 electroporator (NEPA GENE) in a 2 mm gap electroporation cuvette (Bulldog Bio, Cat. #: 12358-346). The settings were: Poring Pulse: 250.0 Volts, 8.0 ms pulse length, 50.0 ms pulse interval, 2 pulses, 10 % decay rate, + polarity; Transfer Pulse: 20.0 Volts, 50.0 ms pulse length), 50.0 ms pulse interval), 10 pulses, 40 % decay rate), +/- polarity.

For recovery, cells were transferred to 10 mL TAP liquid medium plus 80 mM sucrose and incubated with gentle shaking under dim light (<10 μmol photons m^−2^ s^−1^) overnight. Colonies were formed on TAP agar containing 10 μg/ml hygromycin B. Candidate colonies were analyzed by PCR, microscopy, and segregant analysis as shown in Supplementary Figure 1.

CRISPR-Cas9-mediated knockout was performed following a published protocol ^93^, using an sgRNA targeting the *SEP1* region encoding the central GTPase domain: 5’-ACATGTACTTGAGGTCGATGGGG to insert an *AphVII* spectinomycin-resistant cassette.

### Anti-SEP1 antisera

The antisera against SEP1 were produced by Yenzym antibodies, LLC (Brisbane, CA, USA). Synthetic SEP1 peptide C-Ahx-HDGSYTPTEQFRRDPESLS-amide was injected into two rabbits. Both antisera produced a single strong band in Western blotting using whole Chlamydomonas extract. Batch “YZ7085” was used throughout this study.

### Western Blotting

Whole-cell extracts were prepared as described previously^57^. SDS-PAGE was performed using 4-20% Mini-PROTEAN TGX Precast Gels (Bio-Rad, Cat. #: 4561094). After the proteins were transferred onto PVDF membranes, the blots were stained with an anti-SEP1 antiserum (1:2000 dilution, see above) or an anti-AtpB (1:5000 dilution, Agrisera, Cat. #: AS05 085) antibody, followed by an HRP-conjugated anti-rabbit-IgG secondary antibody (1:25000 dilution, Invitrogen, Cat. #: 31466).

### Light microscopy

Imaging of LatB-treated cells was performed using a Zeiss Axio Imager Z2 Widefield Fluorescence Microscope. We used a single Z slice to image cells with a 100X Plan-Apochromat 1.4 NA oil objective. Images were acquired using an Axiocam 506 color camera to visualize natural color of cells.

For live-cell time-lapse imaging, low-melting agarose (LMA) (Bio-Rad, Cat. #: 161-3114) was added to TAP to prepare 1.5% and 2% LMA/TAP media. 1.5% LMA/TAP blocks were prepared by pipetting molten agarose into the cap of a microcentrifuge tube. After solidifying, small blocks were excised from the cap and allowed to dry on a Whatman filter paper for several minutes. Meanwhile, cells were scraped from agar plates using a 10 μl inoculating loop and resuspended in 500 μl liquid TAP media. The cells were pelleted at 2,000 x *g* for 2 minutes. The supernatant was removed and cells were re-suspended in 50 μl of fresh TAP media. A small volume (∼5 μl) of the cell suspension was pipetted onto the 1.5% LMA/TAP blocks, allowed to settle for 5 minutes, and then carefully wicked dry using the torn edge of a piece of Whatman filter paper. The agarose block was then flipped and transferred into an 18-well chambered glass coverslip (Ibidi) and gently pushed down so that the cells sit at the coverslip interface. Afterwards, 2% LMA/TAP was melted, cooled to ∼45°C, and pipetted into the well to seal the agarose block and prevent drying.

Most fluorescence microscopy was performed on a Leica Thunder inverted microscope equipped with an HC PL APO 63X/1.40 N.A. oil-immersion objective lens. Signals were captured using following combinations of excitation and emission filters: 510 nm and 535/15 nm for NG (10%, 300 ms); 550 nm and 595/40 nm for SC (7%, 200 ms); 640 nm and 705/72 nm for chlorophyll AF (1%, 10 ms), with 0.21 μm Z-spacing covering 6-10 μm. All images were processed through Thunder Large Volume Computational Clearing and Deconvolution (Leica). To capture SEP1 localization at alternative stages of the cell cycle, we used a 2-minute interval. To capture SEP1 ring closure we used a minimal acquisition interval of 1 minute 18 seconds. In cases where we wanted to extend time-lapse movies for longer periods of time, we reduced the laser power of NG and SC channels to 5% to minimize photobleaching and phototoxicity. To capture SEP1 localization at iterative chloroplast division sites, we used a 4-minute acquisition interval.

To visualize SEP1 and TOC90/120 colocalization using super-resolution microscopy, we utilized an Andor Dragonfly Spinning Disk Confocal combined with SRRF (Super-Resolution Radial Fluctuations) processing. To visualize NG, samples were excited with a 488-nm diode laser and captured with a 525/50 filter. To visualize SC, samples were excited with a 561-nm diode laser and captured with a 600/50 filter. Samples were imaged with a 63x/1.20 W HC PL APO CORR CS2 (Leica 11506346) water objective. Images were acquired using an Andor Zyla PLUS 4.2 Megapixel sCMOS camera.

### Plasmids

Plasmids used in this study and their constructions are summarized in Supplementary File 1. Plasmid sequence files are available upon request.

### Ultrastructural Expansion Microscopy

Cryo-expansion microscopy was performed as published previously^94^. Briefly, CC-124 WT and *sep1Δ* (*sep1Δ* ARC6-NG-3FLAG) cells were synchronized on agar-TAP plates using a 12 hr:12 hr light:dark cycle (light intensity 60-70 µmol photons m^-2^ sec^-1^) at 25°C. A small amount of cells was transferred in a thin layer to a fresh agar-TAP plate for several consecutive days. 1-3h after beginning of the dark phase, cells were craped from the plate, transferred into TAP medium and centrifuged for 5 min at 500 x *g*. The pellet was resuspended in a smaller volume of TAP. Around 30 µl of cells were put on poly-D- or poly-L-lysine-coated 6-mm round glass coverslips and adhered for at least 10 min.

To fix cells, TAP was removed from the coverslip, residual liquid carefully blotted from the side, and the coverslip was rapidly plunged in liquid ethane using a manual plunge-freezing system (RHOST LLC). The coverslip was transferred quickly into a 1.5-ml tube containing liquid-nitrogen-frozen dry acetone with 0.1% paraformaldehyde (PFA) and 0.02% glutaraldehyde (GA). Immediately afterwards, the tube was placed on dry ice. Coverslips were plunge-frozen successively and incubated overnight in the acetone solution on dry ice in a polystyrene box while shaking.

The next day, dry ice was removed from the box, the lid closed and coverslips incubated for another 45 min while shaking. Next, coverslips were rehydrated in descending percentages of ethanol in ddH_2_O: In a 24-well plate, coverslips were incubated 2x in 100% ethanol with 0.1%PFA and 0.02% GA (cooled to −20°C), 2x 95% ethanol with 0.1% PFA and 0.02% GA (cooled to −20°C), 70% ethanol, 50% ethanol, 25% ethanol, pure ddH_2_O, and then PBS. Incubation was performed for 5 min each. From 70% ethanol onwards, incubation was done at room temperature. After rehydration, coverslips were checked on a light microscope to confirm the side on which the cells were attached before plunging. Coverslips with cells on the bottom were inverted.

Expansion was performed as described previously ^95^. Briefly, fixed cells were incubated in anchoring solution (2% acrylamide, 1.4% formaldehyde in PBS) at 37°C for 3h. For gelation, we worked on a −20°C-cold metal block on ice to slow down the gelation. 5 µl 10% TEMED in nuclease-free water and 5µl 10% APS in nuclease-free water were added to 90 µl monomer solution [23 % (w/v) sodium acrylate, 10 % (w/v) acrylamide, 0.1 % (w/v) N,N’-methylenbisacrylamide in PBS] and vortexed for 2-3 sec. Immediately afterwards, drops of 9 µl were placed on parafilm-coated microscopy slides on the metal block. One by one, coverslips were removed from the anchoring solution, excess liquid removed by blotting from the side, and the coverslips placed on the drops with the cells facing the solution. Coverslips were incubated on the metal block for 5 min, and slides with coverslips carefully transferred to a humid chamber. Gels were polymerized at 37°C for 30 min. Afterwards, coverslips (with gels facing upwards) were incubated in 6-well plates with denaturation buffer (200 mM SDS, 200 mM NaCl, 50 mM Tris in ddH_2_O, pH 9.0) while shaking. Once the gels detached from the coverslips (after 20-30min), they were transferred into 1.5-ml tubes filled with fresh denaturation buffer and incubated at 95°C for 90min in a thermoblock. Next, gels were transferred into MilliQ water in wells of a 6-well plate. For full expansion, gels were incubated 3x for at least 30 min in MilliQ water.

For immunostaining, MilliQ water was exchanged for PBS (usually 2×10min with shaking) to shrink gels and fit them into wells of a 12-well plate. Cells were blocked with filtered 3% BSA/0.1% Tween-20 in PBS for 30 min at 37°C, shaking. Next, cells were washed once in 0.1% Tween-20 in PBS for 10 min at RT, shaking, and subsequently incubated with primary antibodies in antibody solution (filtered 1% BSA, 0.05% Tween-20 in PBS) for 2.5h at 37°C, shaking. The following primary antibodies were used: guinea pig anti-alpha- and anti-beta tubulin monobodies (ABCD-Antibodies, AA345 and AA344, both 1:500 dilution) and rabbit anti-CrSEP1 serum (see above). After washing gels with 0.1% Tween-20 in PBS for 3×10min at RT, shaking, cells were incubated with secondary antibodies and DAPI (1:500 dilution) in antibody solution (filtered 1% BSA, 0.05% Tween-20 in PBS) for 2.5h at 37°C, shaking. The following secondary antibodies were used: goat anti-guinea pig Alexa Fluor 568 (Invitrogen, Cat. #: A-11075, 1:1000 dilution) and donkey anti-rabbit Alexa Fluor 647 (Invitrogen, Cat. #: A-31573, 1:1000 dilution). Gels were washed 3×10min with 0.1% Tween-20 in PBS at RT, shaking, and washed 1×10min with PBS. Next, gels were incubated with Alexa Fluor 488 NHS Ester (ThermoFisher, Cat. #: A20000, 2 µg/ml final concentration) in PBS overnight at 4°C, shaking, followed by 3×10min washes with 0.1% Tween-20 in PBS at RT, shaking. For full expansion of the gels, they were placed in MilliQ water for three times for at least 30 min.

For imaging, the diameter of the fully expanded gels was measured using a ruler. The expansion factor (around 4-fold) was determined by dividing the diameter of the gels by the diameter of the coverslip (6 mm). The surface of the gels was carefully dried with a tissue to identify the side of the gel which contains the cells (the other side shows the typical pattern from the parafilm during gelation). Gels were cut and mounted with cells facing down on poly-L- or poly-D-lysine-coated 2-well Ibidi dishes with glass bottom. Wells were filled with ddH_2_O to avoid drying/shrinkage of the gels during imaging. Cells were imaged using a Zeiss LSM980 Airy Fast Microscope in the 8-fold multiplexing super-resolution mode. Images were acquired using a C-Apochromat 40x/1.2NA water immersion objective, diode- and diode-pumped solid-state lasers and respective emission filters. To acquire Airyscan super-resolution images, the pixel size was set to 0.04 µm in x-y. A 3x or 4x zoom was used to image the cells, resulting in a total image size of 70.7 x 70.7 µm (1716 x 1716 pixels) and 53×53 µm (1288 x 1288 pixels), respectively. The pixel dwell time was between 0.61 and 0.81 µs, respectively. Laser powers and z-stack sizes (with constant 0.16 µm Z-intervals) were adjusted based on the staining and cell size, respectively. Scale bars in images were adjusted for the expansion factor.

### Angular Integration Analysis of Chloroplast Division Proteins

Cells were synchronized as previously described ^55^ using a 12 h:12 h light:dark chamber at 25°C. After three days of continuous dilutions and synchronization, the cells were transferred onto TAP + 3 μM LatB (or TAP + 0.1% DMSO control) 10 minutes prior to the start of the dark cycle. After being kept in the chamber for another 2 hours, cells were collected and imaged on 1.5%LMA/TAP pads containing either 0.1% DMSO or 3 μM LatB. Imaging was performed on two separate days, and data from both days were pooled, yielding 20-30 cells across 6-7 fields for analysis.

For analysis, although our cultures were synchronized, natural variation in cell cycle progression resulted in cells at different cell cycle stages. It was important to select cells of similar sizes during division. Only cells that had not yet divided their chloroplasts were analyzed. To eliminate bias, individual cells were solely selected based on the chlorophyll autofluorescence channel. Chloroplasts during division exhibit a distinct morphology: a widening of the chloroplasts and shortening of the chloroplast lobes (cup-like to bowl-like transition). After selection, images of cells were rotated to approximate the AP-axis based on chloroplast geometry. These cells were then average Z-projected. Circular ROI was generated for each cell, and coordinates were recorded. These coordinates were then used in a custom script to measure the signal intensity using a polar coordinate system, obtaining measurements for all channels as a function of theta.

### FLAG Immunoprecipitation in *C. reinhardtii*

About 4 grams of harvested cell material was resuspended in cold 2× IP buffer at a 1:1 weight-to-volume ratio of cold 2X IP buffer [100 mM HEPES pH 6.8, 100 mM KOAc, 4 mM Mg(OAc)_2_, 2 mM CaCl_2_, 400 mM sorbitol], freshly supplemented with protease inhibitor cocktail (Roche, Cat. #: 11873580001), and phosphatase inhibitor cocktail (Roche, Cat. #: 04906837001). The cell suspension was then snap-frozen in liquid nitrogen. Frozen pellets were ground using a cryomill (Retsch MM400, Cat. #: 70354). Cell powder was then allowed to slowly thaw on ice for ∼45 minutes. Once defrosted, the sample was transferred to a Kontes Duall #21 homogenizer and dounced 20 times. 500 μl samples were transferred to pre-chilled microcentrifuge tubes. An equal volume of cold 1X IP buffer + 2% digitonin was added (final digitonin concentration 1%). Samples were incubated at 4°C for 40 minutes with nutation. Lysates were clarified by spinning at 4°C for 30 minutes at full speed in a tabletop centrifuge.

Lysates were incubated with magnetic Dynabeads protein G conjugated to anti-FLAG M2 antibody (Sigma, Cat. #: F1804), previously washed with 1X IP buffer and equilibrated in 1X IP buffer + 0.1% digitonin, at 4°C for 90 minutes on a rotating platform. Samples were then washed 4 times with 1X IP buffer + 0.1% digitonin, and then eluted with 1X IP Buffer + 2 μg/μl 3xFLAG peptide by gentle shaking at 4°C for 30 minutes.

### Yeast Two Hybrid

Yeast two-hybrid assays were performed as described previously^96^. Briefly, the *S. cerevisiae* reporter strain PJ69-4A was cotransformed with a plasmid containing GAL4 activation domain fused to the SEP1 GTPase domain and a second plasmid containing GAL4 DNA-binding domain fused to GTPase domains of SEP1, TOC34, TOC90, and TOC120.

Colonies were selected for both leucine and tryptophan auxotrophy to confirm transformation with both plasmids. To test the GAL4-driven transcription of the *HIS3* reporter, transformants were plated on synthetic complete media minus leucine, tryptophan, and histidine, supplemented with 1 mM 3-amino-1,2,4-triazole.

### SEP1-BioIDG3 proximity labeling

WT CC-124 and strains expressing SEP1–BioIDG3–NeonGreen were streaked onto fresh TAP plates and placed under a 12-hr:12-hr dark:light cycle at 21°C. The cells were synchronized for two days and then transferred to TAP agar plates supplemented with 1 mM biotin for an additional 24 hrs, for 12 hrs in dark at 21°C and 12 hrs in light at 33°C. Portions of cells were harvested at defined time points during the light phase (10, 11, and 12 hrs) by scraping into TAP medium, washing twice with fresh TAP medium to remove excess biotin, and then pelleting by centrifugation. The cell pellets (∼0.3 grams wet weight, approximately 1.5 x 10^8^ cells) from three time points were pooled and resuspended in 1.5 mL RIPA-based lysis buffer [25 mM Tris–HCl pH 7.4, 300 mM NaCl, 1 mM DTT, 5 mM MgCl2, 0.1 mM PMSF, 1X EDTA-free HALT protease inhibitor (ThermoFisher, Cat. #: 78425), 0.1% (w/v) SDS, 0.5% (w/v) deoxycholic acid, and 1% (v/v) Triton X-100)], and snap-frozen in liquid nitrogen. Lysis was performed by five cycles of freezing at −80°C for 60 min followed by thawing at room temperature (15-25 min per cycle). Prior to streptavidin affinity pulldown, free biotin was removed with a Zeba Spin Desalting Column (ThermoFisher, Cat. #: 89891). Pierce Streptavidin Magnetic Beads (ThermoFisher, Cat. #: 88816) were washed three times with five volumes of RIPA-based lysis buffer prior to use. 4.2 grams of precleared lysates were incubated with 300 µg magnetic beads overnight at 4°C with gentle rotation. Beads were washed three times with 150 µL cold RIPA-based lysis buffer, followed by brief incubation with 150 µL of cold 1 M KCl. Beads were subsequently washed twice with RIPA-based lysis buffer. Bound proteins were eluted at 90°C for 8 min in 40 µL elution buffer (25 mM Tris pH 7.5, 2% SDS, 10 mM DTT, 2 mM biotin). The eluents were frozen until submitted for mass spectrometry.

### LC-MS/MS proteomics sample processing, data collection and data analysis

Samples were brought to 5% SDS and reduced with 10 mM DTT for 20 min at 32°C, alkylated with 25 mM iodoacetamide for 30 min at room temperature, then supplemented with a final concentration of 1.2% phosphoric acid and 200 µL of S-Trap (Protifi) binding buffer (90% MeOH/100mM TEAB). Proteins were trapped on the S-Trap micro cartridge, digested using 1 µg of sequencing grade trypsin (Promega) for 1 hr at 47°C, and eluted using 50 mM TEAB, followed by 0.2% FA, and lastly using 50% ACN/0.2% FA. All samples were then lyophilized to dryness. Samples were resuspended in 40 µL of 1% TFA/2% acetonitrile with 12.5 fmol/µL of yeast ADH. A study pool QC (SPQC) was created by combining equal volumes of each sample and run periodically throughout the study.

LC/MS/MS was performed using an EvoSep One UPLC coupled to a Thermo Orbitrap Astral high resolution accurate mass tandem mass spectrometer (Thermo). Briefly, 30% of each sample was loaded onto an EvoTip which was eluted onto a 1.5 µm EvoSep 150 µm ID x 15 cm performance (EvoSep) column using the SPD30 gradient at 55°C. Data collection on the Orbitrap Astral mass spectrometer was performed in a data-independent acquisition (DIA) mode of acquisition with a r=240,000 (@ m/z 200) full MS scan from m/z 380-980 in the OT with a target AGC value of 4e5 ions. Fixed DIA windows of 4 m/z from m/z 380 to 980 DIA MS/MS scans were acquired in the Astral with a target AGC value of 5e4 and max fill time of 6 ms. HCD collision energy setting of 27% was used for all MS2 scans. Data were imported into Spectronaut v20 (Biognosis) and MS/MS data was searched against a Trembl *C. reinhardtii* database (downloaded in 2024), a common contaminant/spiked protein database (bovine albumin, bovine casein, yeast ADH, etc.), and an equal number of reversed-sequence “decoys” for false discovery rate determination. Database search parameters included fixed modification on Cys (carbamidomethyl) with variable modification on Met (oxidation). Full trypsin enzyme rules were used along with 10 ppm mass tolerances on precursor ions and 20 ppm on product ion. Spectral annotation was set at a maximum 1% peptide false discovery rate based on q-value calculations. Peptide homology was addressed using razor rules in which a peptide matched to multiple different proteins was exclusively assigned to the protein has more identified peptides.

Relative abundances of annotated precursor ions were measured based on MS2 fragment ions of selected ion chromatograms of the aligned features across all runs. Next, signals corresponding to yeast BSA, trypsin, bovine casein, and keratin were temporarily excluded, and remaining precursors were subjected to a trim mean normalization in which the highest and lowest 10% of the signals were excluded and the remaining mean value was used to normalize precursor values across all samples. Missing values were then imputed at the lowest 2% of the measured distribution of intensities of all precursors. Finally, the IQ_MaxLFQ ^97^ rollup strategy was deployed to convert from the precursor level to the protein level based on all remaining precursor values.

### cTP-SC Reporter Assays

Strains expressing cTP reporter constructs for FTSZ1, FTSZ2, ARC6, and RBCS2 in wild-type and *sep1Δ* genetic backgrounds were grown for 4 days on TAP agar with daily refreshing. Imaging was performed on two independent days and data from both days were pooled, yielding approximately 500 cells total across 4–5 fields for analysis. For imaging, unsynchronized cells were used. Cells were transferred using a sterile loop and spread as a thin layer onto 1.5% LMA/TAP pads.

Images were acquired with 0.4 µm Z stack spacing spanning 8 µm. Illumination and acquisition settings were as follows: DIC: 10 ms exposure, mScarlet (SC channel): 550 nm excitation, 40% laser power, 300 ms exposure, Chlorophyll fluorescence (Chl. channel): 640 nm excitation, 2% laser power, 20 ms exposure. All images were processed using Leica THUNDER Large Volume Computational Clearing and Deconvolution.

All subsequent image processing and quantification were performed in Fiji/ImageJ using the medial optical section of each Z-stack. The Chl. channel was used to create the chloroplast mask using “Auto Threshold: Default (IsoData).” To create a whole-cell mask, the chlorophyll channel was thresholded using Huang’s fuzzy thresholding method^98^, which emphasizes overall object boundaries rather than internal signal structure. Because Chl. is often reduced in the pyrenoid region, a “hole” is often created inside the chloroplast mask. These “holes” were manually filled prior to downstream analysis. The ratio of reporter signals between the chloroplast mask and the whole cell (which contains the chloroplast mask) was calculated for each cell.

For cycloheximide treatment experiment, wild-type cells expressing cTP^RBCS2^-SC were grown unsynchronized in liquid TAP for 2 days at room temperature under continuous illumination at 50 µmol photons m⁻² s⁻¹ in 6-well plates. Cultures were refreshed the day prior to imaging.

Cycloheximide (Sigma-Aldrich, Cat. #: 01810-5G) was added to a final concentration of 30 µg/mL to inhibit cytosolic translation. Cells were imaged at 0, 15, 30, 60, 150, 360, and 480 minutes following cycloheximide addition. At each time point, an aliquot of culture was transferred to a 1.5 mL microcentrifuge tube and briefly centrifuged to concentrate cells. Two microliters of the concentrated cell suspension were placed onto an agar pad for imaging. Imaging analysis was performed as described above.

### Septin & TOC Phylogenetic Analysis

*Arabidopsis thaliana* and *C. reinhardtii* TOC GTPase protein sequences were obtained from TAIR and Phytozome, respectively. To identify TOC33-family homologs in early diverging Archaeplastida, we first performed a BLASTP search against the rhodophyte *Porphyridium purpureum*. The resulting hit was then used in iterative BLASTP searches to identify representative TOC33-family proteins from Rhodophyta and Glaucophyta. Representative eukaryotic septin sequences were obtained from previous publications.

The GTPase domain of SEP1 was first defined based on the crystal structure reported by Pinto et al. (2017). A BLASTP database was generated from the collected TOC and septin sequences, and the SEP1 GTPase domain was used as the query. Based on these BLASTP results, the corresponding GTPase domains of all septin and TOC proteins were defined. These GTPase domain sequences were aligned using Anchored DIALIGN. As an anchor, we specified a single conserved arginine residue shared between *C. reinhardtii* SEP1 (R239) and *C. reinhardtii* TOC34 (R198), corresponding to *A. thaliana* TOC33 R130.

The resulting multiple sequence alignment was used to infer a maximum-likelihood phylogeny in IQ-TREE under the LG+I+G4 substitution model with 1,000 ultrafast bootstrap replicates. Model selection was performed using Bayesian information criterion (BIC) scores.

### Heterologous Expression Experiments

#### Volvox carteri

The recipient strain TNit-1013 was used for transformation; it originates from wild-type strain HK10 (aka Eve10) ^99,100^. Algae were cultured in standard *Volvox* medium ^101^. Light and temperature regime for cultivation were as described previously ^102,103^. For subcellular localization of VcSEP1 in *V. carteri*, vector pBlue-VcSEP1-YFP was constructed, allowing for expression of VcSEP1 combined with a C-terminal YFP fluorescence tag, following the strategy described previously ^104^. Stable nuclear transformation was achieved by particle bombardment performed as described elsewhere with minor modifications ^105,106^. pBlue-*Vc*SEP1-YFP was cotransformed with plasmid p*Vc*NR15 to complement the nitrate reductase gene ^107^. Selection followed previously described procedures based on the algae’s capability to use nitrate as nitrogen source ^108^. Genotyping was carried out analogously to von der Heyde et al., 2015 and proved that both plasmids were integrated in all selected strains (12 strains in total, 4 independent). Subsequent fluorescence screening identified three independent strains with detectable YFP signal.

Sample preparation for live cell imaging was performed as previously described ^109^. For Imaging of *Vc*SEP1-YFP, a LSM780 confocal laser scanning microscope equipped with LCI Plan Neofluar 63x/NA 1.3 and Plan-Apochromat 10x/0.45 M27 objectives was used (Carl Zeiss GmbH, Oberkochen, Germany). An argon-ion laser was used for excitation at 514 nm. Fluorescence was detected in two channels simultaneously: YFP at 517-553 nm and chlorophyll at 650-700 nm. Images were recorded with 12-bit depth using the ZEN black imaging software (ZEN 2011, Carl Zeiss GmbH, Germany). Fiji was used for image processing and analysis.

#### Physcomitrium patens

Moss protoplasts were transformed with 30 µg of pTKUbi-GFP-SEP1 plasmids using PEG mediated protoplast transformation as previously described ^110^. Protoplasts were incubated for three days in Hoagland’s medium [4mM KNO_3_, 2mM KH_2_PO_4_, 1mM Ca(NO_3_)_2_, 89µM Fe citrate, 300µM MgSO_4_, 9.93µM H_3_BO_3_, 220nM CuSO_4_, 1.966µM MnCl_2_, 231nM CoCl_2_, 191nM ZnSO4, 169nM KI, 103nM Na_2_MoO_4_, and 1% sucrose] supplemented with 8.5% mannitol and 10 mM CaCl_2_. Three days after transformation, half of the protoplasts were collected by centrifugation at 250 x *g* and then resuspended in 100 µL of the same solution described above. Protoplasts were placed on a glass-bottom dish and imaged using a laser-scanning confocal microscope. The other half of the transformation was plated and allowed to regenerate into individual plants. Stable transgenic lines were selected by the presence of GFP fluorescence.

Transgenic plants expressing GFP-SEP1 were propagated and protonemata were placed on an agar pad for confocal imaging as previously described ^111^. Images were acquired on a Nikon A1R confocal microscope system with a 1.49 NA 60X TIRF oil immersion objective (Nikon) at room temperature. Image acquisition was controlled by NIS-Element AR 4.1 software (Nikon). Laser illumination at 488 nm was used for mEGFP and chloroplast auto-fluorescence (laser power, 0.5–1%; Gain, 40–60; PMT offset, 28–33). Emission filters were 525/50 nm for mEGFP and 600/50 nm for chloroplast auto-fluorescence.

#### Arabidopsis thaliana

*A. thaliana* Col-0 were transformed using the floral dip method with plasmid pSD045 ^112^. Transformed seeds were selected based on sulfadiazine resistance. Selected T2 seedlings (4 days old, grown in the dark) were analyzed by confocal laser scanning microscopy using a Zeiss LSM880 with Airyscan. Excitation/emission wavelengths were 488 nm/492-570 nm for GFP and 561 nm/588-696 nm for chlorophyll fluorescence.

For *A. thaliana* protoplast expression, the SEP1 coding sequence was fused in-frame with YFP and cloned into the pUC19 plasmid. The protoplasts were prepared as described previously ^113^. Briefly, *A. thaliana* leaves were harvested at 3.5 weeks and digested to isolate protoplasts. A 150 µL aliquot of the protoplast suspension was transfected with 10 µg plasmid DNA purified using ZymoPURE II Plasmid Maxiprep Kit (ZYMO Research), followed by overnight incubation at 23°C.

#### Nicotiana benthamiana

For transient expression in *N. benthamiana*, an overnight culture of *Agrobacterium* strain GV3101 carrying SEP1-YFP in the pCambia1300 backbone was collected by centrifuging at 6000 rpm for 3 min and resuspended in infiltration buffer containing 10 mM MES, 10 mM MgCl_2_, and 200 µM Acetosyringone. The suspension was adjusted to OD_600_ = 0.6 and infiltrated into fully expanded *N. benthamiana* leaves. Leaf discs were collected after 24-hr inoculation, and the confocal microscopy images were aquired using the same parameters as described previously_114._

For immunoprecipitation mass spectrometry experiments, transient expression of SEP1-YFP in *N. benthamiana* was performed as described above using an *Agrobacterium* suspension at OD_600_ = 0.2. 3 grams of infiltrated leaf tissue was collected and immediately frozen in liquid nitrogen. Total proteins were extracted using prechilled 2.5X IP buffer containing 10% glycerol, 25 mM Tris-HCl, pH 7.5, 1 mM EDTA, 150 mM NaCl, 10 mM DTT, 1× protease inhibitor cocktail (Roche, Cat. #: 04906837001), and 0.5% NP-40. Lysates were centrifuged at 12,000 rpm for 10 min at 4°C twice, and the supernatant was collected and incubated with 30 µL GFP-trap Agarose (ChromoTek, Cat. #: gtma) beads at 4°C overnight with gentle rotation. The beads were then collected using a magnetic rack and washed four times with ice-cold IP buffer at 4°C and the beads were subjected to LCMS analysis.

### Phylogenetic Identification of TOC homologs in *N. bethanamiana*

Version 1.1 of the *N. benthamiana* proteome and annotations were downloaded from NbenBase. *A. thaliana* TOC33- and TOC159-family GTPases as query, BLASTP searches were performed locally to identify additional paralogs. Paralogs were further validated by a reciprocal universal best hit approach against the *A. thaliana* proteome. The candidate list of *N. benthamiana* TOC GTPases were then collected into a list of TOC GTPases including those from *A. thaliana* and *C. reinhardtii*. These TOC sequences were then used to generate a maximum likelihood phylogenetic tree on IQTREE with an 0.pfam+G4 model according to the BIC score.

## ACKNOWLEDGMENTS

We thank Erik Soderblom, Greg Waitt and Tricia Ho of the Duke University School of Medicine Proteomics and Metabolomics facility for their help with LC-MS analysis of SEP1-BioIDG3 and heterologous SEP1 mass spectrometry in tobacco, the Mass Spectrometry and BioOptics facilities of The Vienna BioCenter Core Facilities for Chlamydomonas IP-MS, Yasheng Gao and Lisa Cameron of the Duke Light Microscopy Core Facility for assistance with super resolution live cell imaging, Yixuan Chen in the Dong lab for assistance with confocal microscopy of tobacco, the Light Microscopy Technology Platform (LiMiTec) for allocation of confocal microscopy facilities at Bielefeld University, Kordula Puls for technical assistance, Xinnian Dong for her advice on *Arabidopsis* and *Nicotiana* experiments, Yaning Yuan in the Onishi lab for assistance with SEP1-BioIDG3 data analysis, and Andrew Countryman for help with image analysis. We thank John Pringle for gift of yeast two-hybrid strains and Felix Kessler for gift of yeast two-hybrid vector plasmids. This work was supported by National Science Foundation Grant MCB #1818383 to John R. Pringle and M.O., National Science Foundation CAREER #2337141 to M.O., Duke University Trinity College of Arts & Sciences, the Austrian Academy of Sciences and the Austrian Science Fund #AST1678424 to S.R., EMBL core support to C.S.S., N.B., and G.D., European Commission (ERC, KaryodynEVO, 101078291) to C.S.S. and G.D. G.D. additionally acknowledges the support of the EMBO Young Investigator Programme. This project has received funding from the European Union’s Horizon Europe research and innovation programme under the Marie Skłodowska-Curie grant agreement No 101109569. Financial support for the work on *V. carteri* was provided by the Bielefeld Young Researchers’ Fund awarded to E.L.v.d.H. Support for S.D. was partially provided by the National Health Institute T32GM142605 training grant to the Duke Program in Cell and Molecular Biology. S-A.C. was a recipient of the Hung Taiwan-Duke University Fellowship. D.S. was a recipient of the Duke Biological Sciences Undergraduate Research Fellowship.

---

**Supplementary Figure 1.**
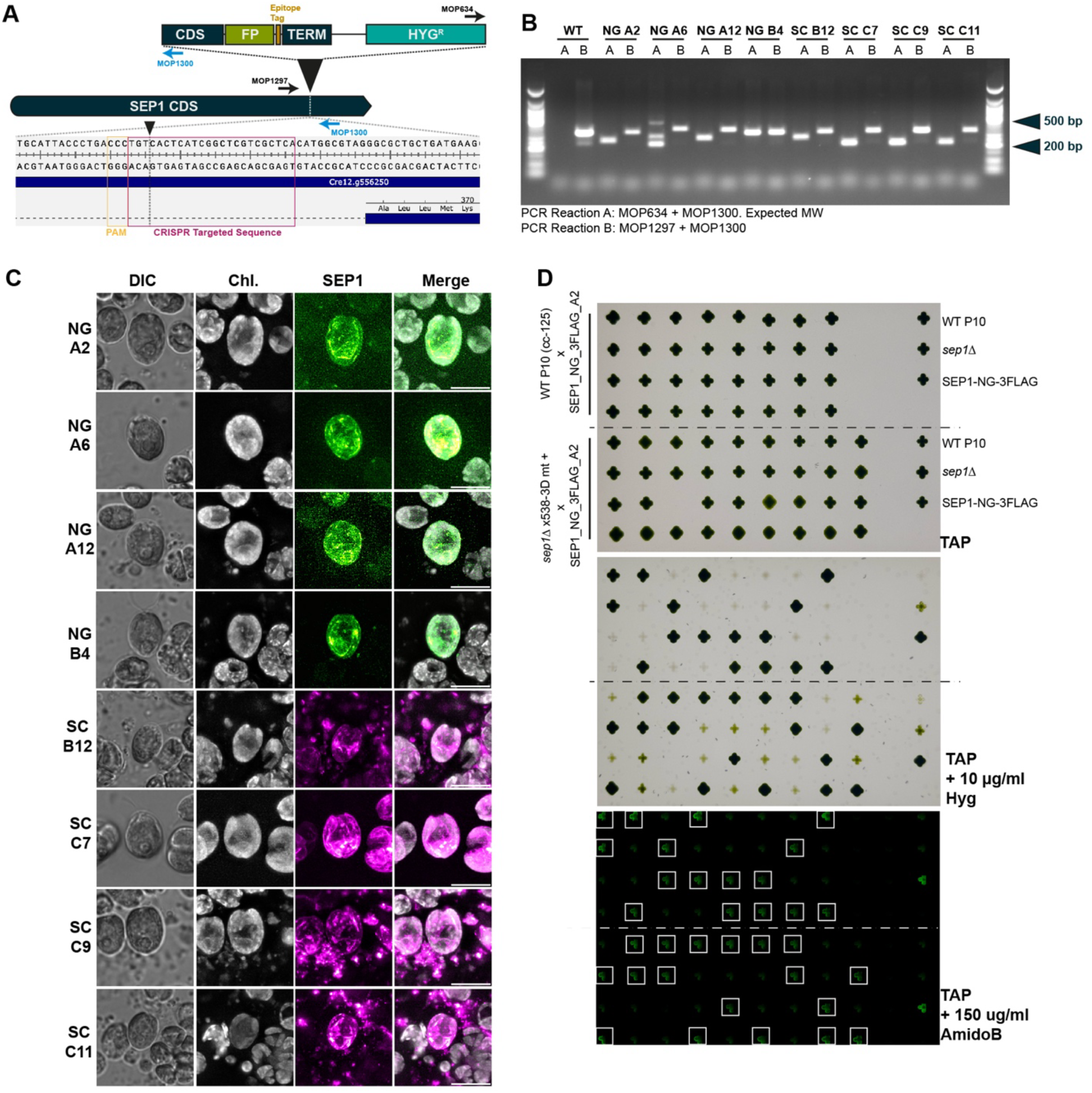
Validation of endogenously tagged SEP1. (A) Cartoon schematic of CRISPR-mediated donor DNA insertion into the *SEP1* genomic locus. (B) Colony PCR of listed reactions utilizing primers annotated in (A). (C) Fluorescence microscopy of individual clones. Max Z-projections. Scale bar: 10 μm. (D) Genetic linkage analysis of SEP1-NG-3FLAG clone A2 when crossed with WT and *sep1Δ* lines. Colonies were plated on TAP, TAP + 10 μg/ml hygromycin, and TAP + 150 μg/ml amido black (imaged using YFP filters).

**Supplementary Figure 2.**
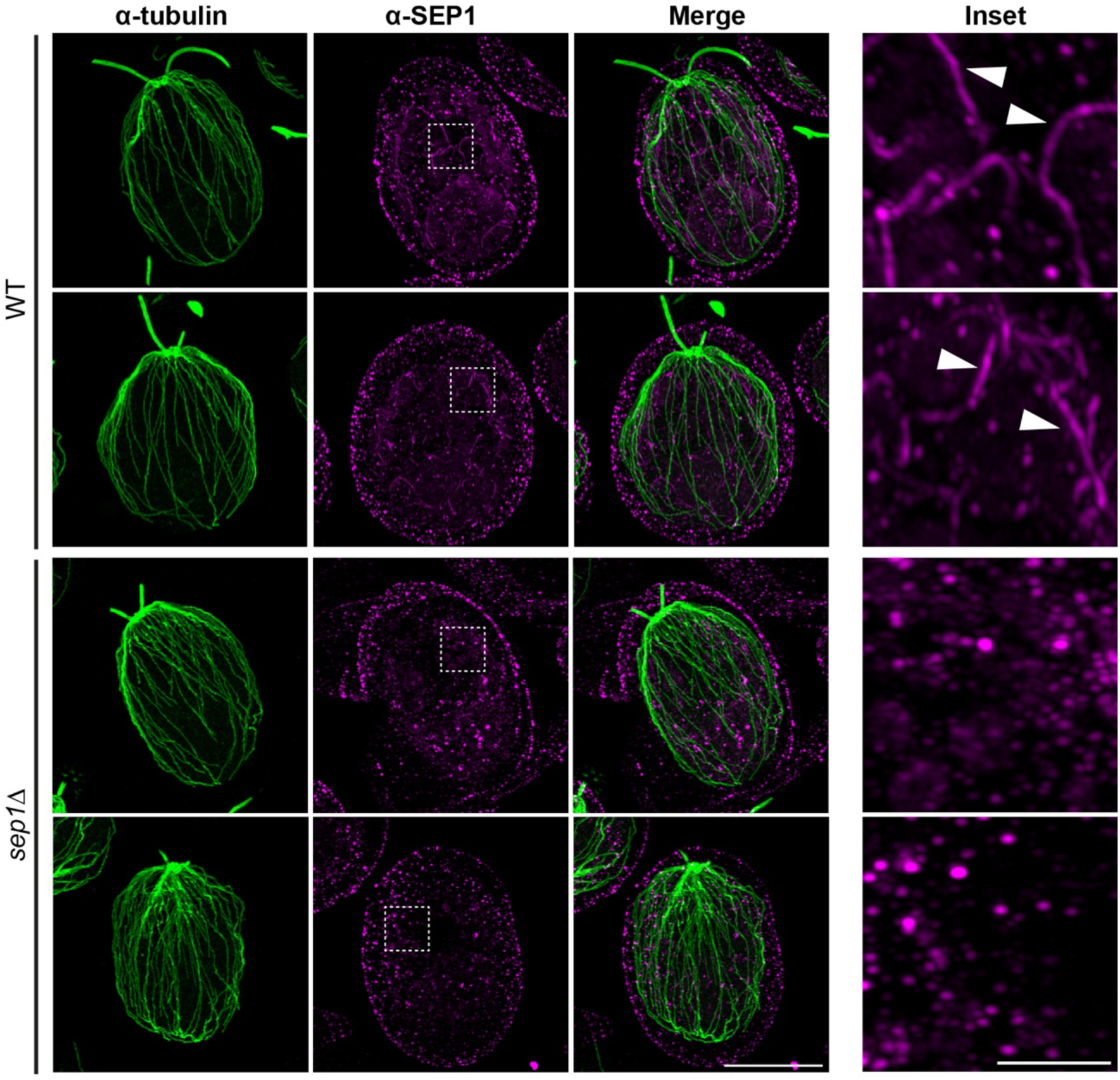
Validation of U-ExM SEP1 signal. Staining of tubulin and SEP1 in WT and *sep1Δ* cells. Scale bar, 5 μm (adjusted for expansion). SEP1 filaments are indicated with arrowheads in WT sample (inset). No filaments were observed in *sep1Δ* cells. Scale bar, 1 μm (adjusted for expansion).

**Supplementary Figure 3.**
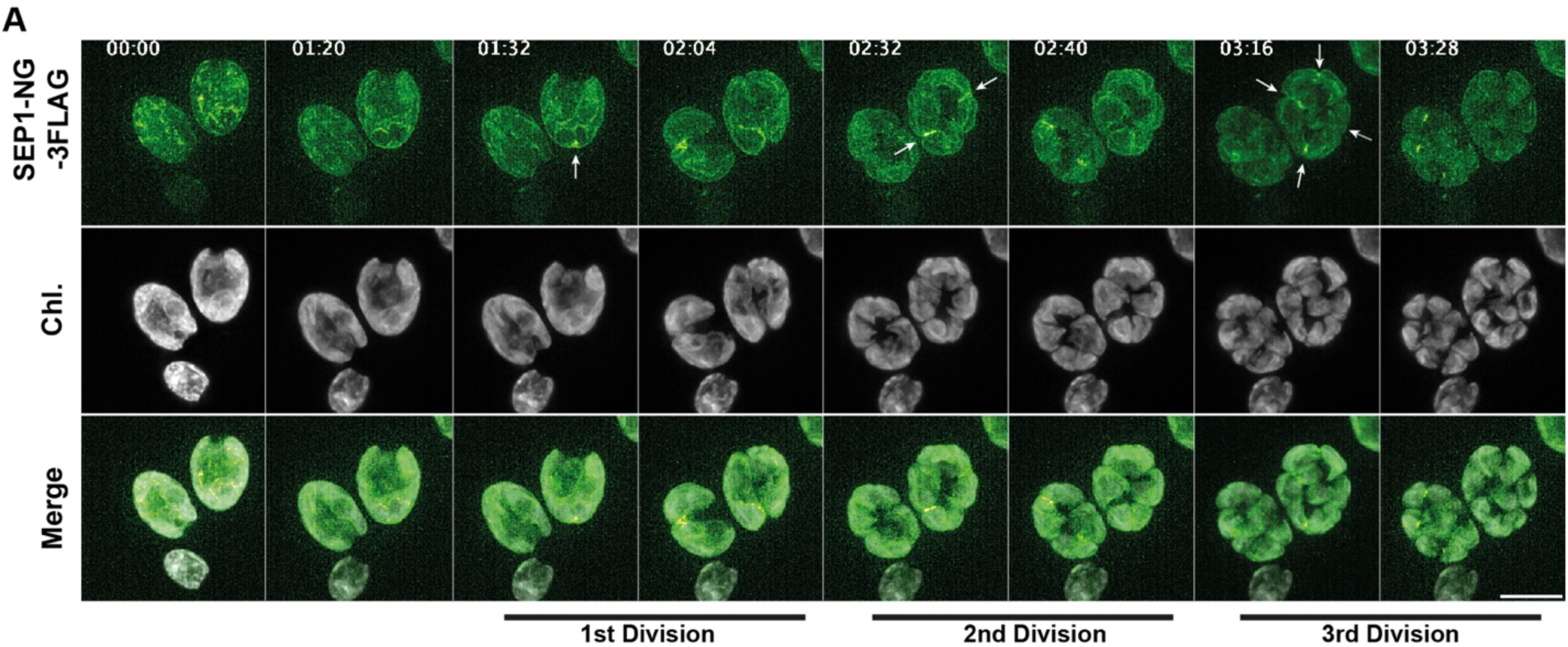
Extended time-lapse series of SEP1 localization during iterative rounds of cytokinesis. SEP1-NG-3FLAG enrichments at iterative chloroplast division sites are indicated with arrows. Time in hh:mm. Scale bar: 10 μm.

**Supplementary Figure 4.**
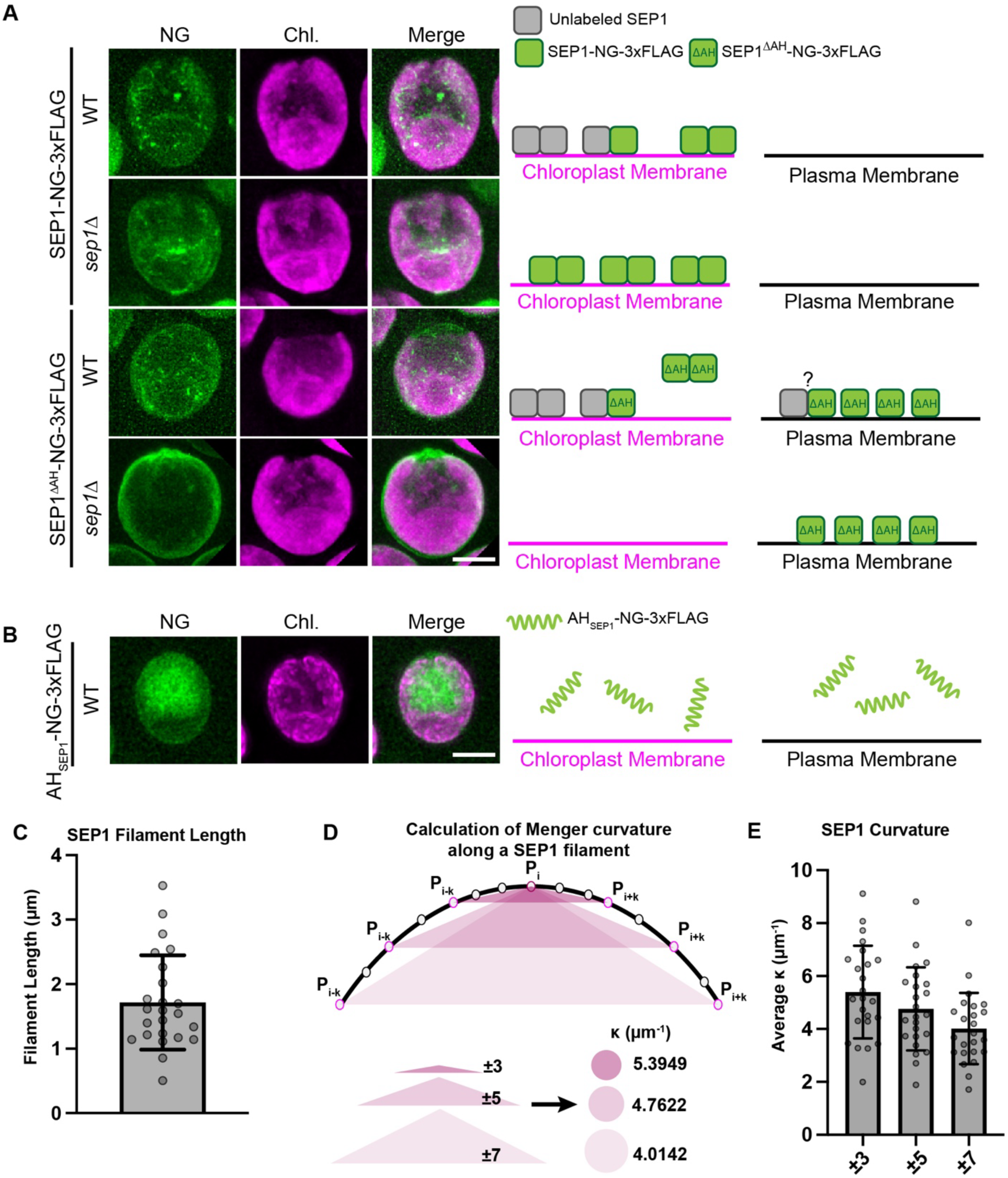
**SEP1 contains a chloroplast targeting amphipathic helix. (**A) Localization of SEP1-NG-3FLAG and SEP1^ΔAH^-NG-3FLAG in WT and *sep1Δ* cells. Scale bar, 10 μm. (B) Localization of the SEP1 AH domain alone (AH_SEP1_-NG-3FLAG) in WT cells. Scale bar, 10 μm. (C) Measurement of individual SEP1 filament lengths using BigTrace plugin in Fiji. μ = 1.717 μm. σ = 0.7329. n = 24. (D) Description of calculation of Menger curvature along a SEP1 filament. Curvature at each point along the filament was calculated using the Menger curvature formula based on triplets of points spaced symmetrically around the point of interest. Specifically, for a point P_i_, curvature was computed using the triangles formed by (P_i-k,_ P_i,_ P_i+k_) with k = 3, 5, and 7, corresponding to increasing window sizes of adjacent points along the filament. (E) Average curvature along filaments. ±3: μ = 5.395, σ = 1.752. ±5: μ = 4.762, α = 1.571 ±7: μ = 4.014, α = 1.346.

**Supplementary Figure 5.**
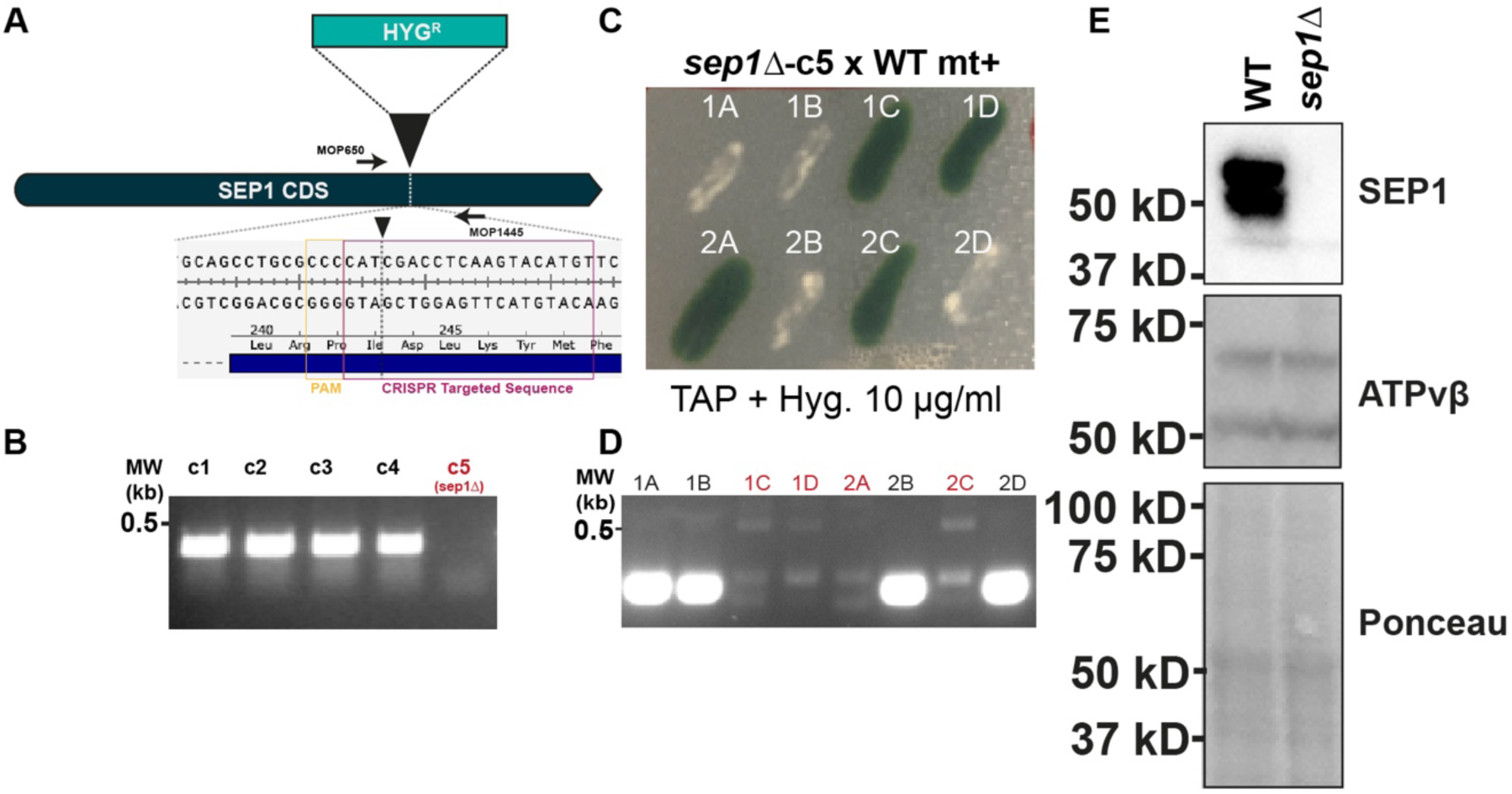
Validation of CRISPR-generated *sep1Δ* strain. (A) Cartoon schematic of CRISPR-mediated insertion of hygromycin-resistance cassette (*HYG^R^*) to disrupt *SEP1*. (B) PCR confirmation of disrupted *SEP1* genomic locus. A *sep1Δ* clone is indicated in red. (C) Genetic linkage analysis confirms the insertion of a single resistance marker cassette. (D) Colony PCR of segregants in (C) confirming a linkage between the *sep1Δ* allele and hygromycin resistance. (E) Western blotting using SEP1 antibody confirms loss of SEP1 protein.

**Supplementary Figure 6.**
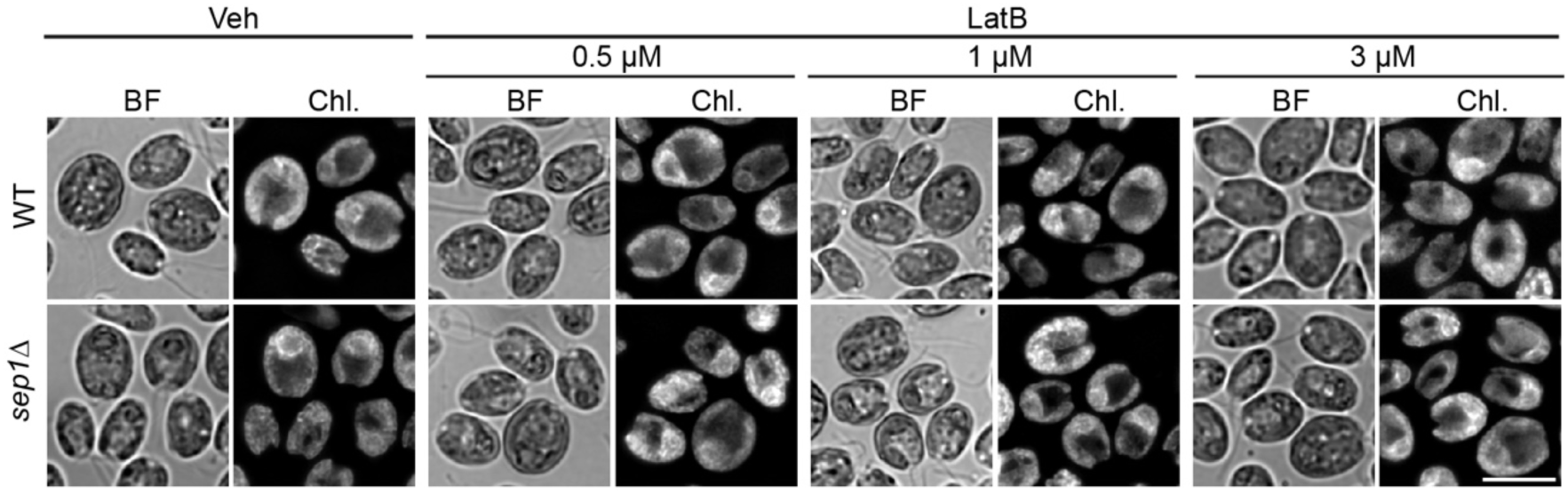
Loss of *SEP1* alone does not disrupt cell or chloroplast morphology. WT and *sep1Δ* cells were treated either with 0.1% DMSO (Veh) or a range of LatB concentrations (0.5 μM, 1 μM, and 3 μM). Scale bar, 10 μm.

**Supplementary Figure 7.**
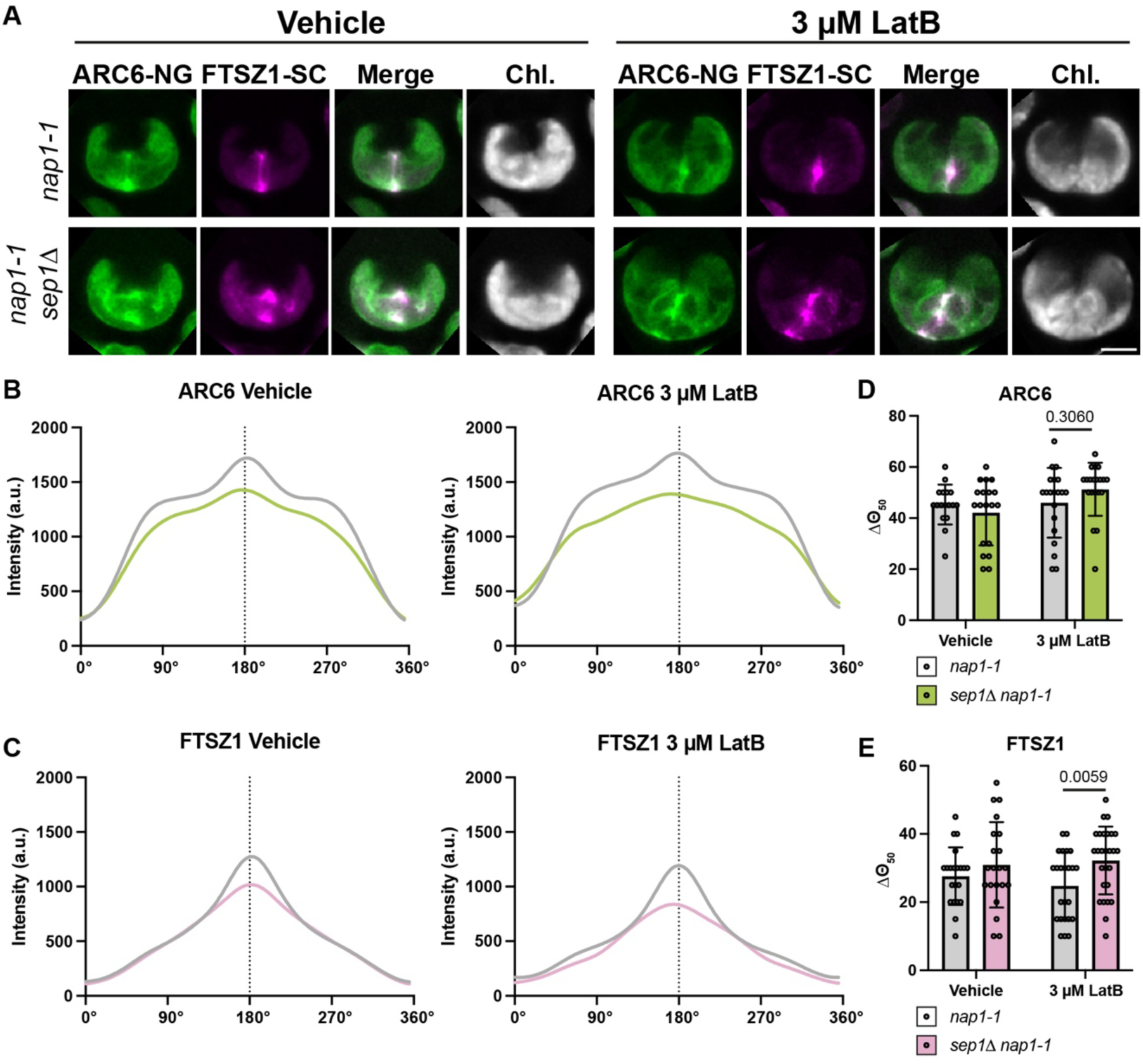
Loss of *SEP1* impairs spatial patterning of FTSZ1. (A) ARC6-NG-3FLAG and FTSZ1-SC-PA localization in *nap1-1* and *sep1Δ nap1-1* cells treated with and without LatB. Vehicle, 0.1% DMSO. Scale bar, 5 μm. (B) Polar plots of ARC6-NG-3FLAG intensity distribution in untreated and LatB-treated cells. (C) Polar plots of FTSZ1-SC-PA intensity distribution in untreated and LatB-treated cells (D) Mean Δθ_50_ values from individual traces in (B). (E) Mean Δθ_50_ values from individual traces in (C). Group comparisons were assessed using one-way ANOVA followed by individual Šidák pairwise comparison.

**Supplementary Figure 8.**
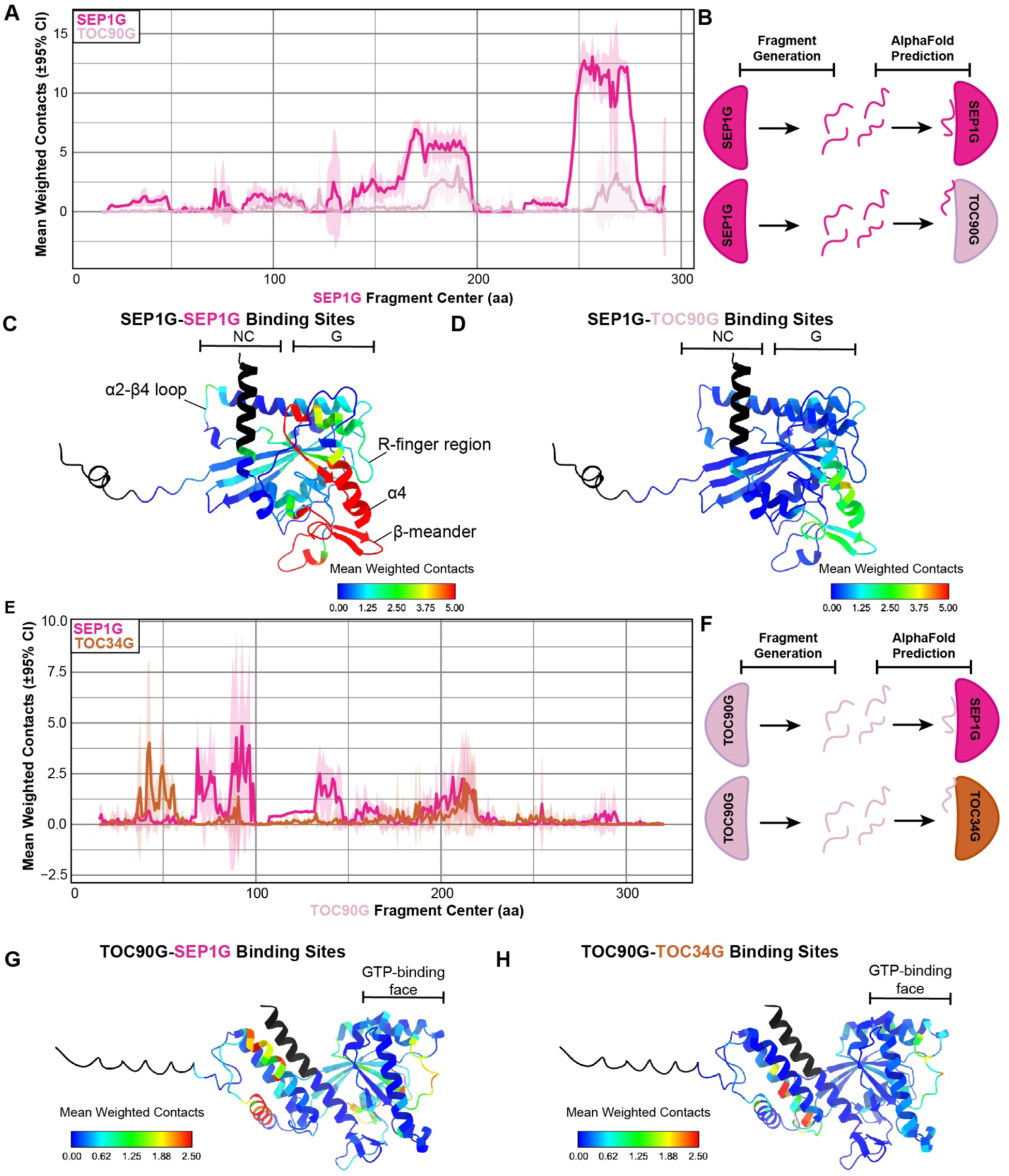
FragFold predictions of SEP1-TOC90 binding sites relative to TOC34-TOC90 binding sites. (A) FragFold predicted mean weighted contact score for SEP1G homodimer and SEP1G-TOC90G heterodimer. (B) Cartoon schematic of FragFold workflow used in (A). (C-D) Predicted SEP1G-homodimer (C) and SEP1G-TOC90G (D) contact sites on SEP1G monomer. NC- and G-interfaces are indicated. Color scaled by mean weighted contact score. (E) FragFold predicted mean weighted contact score for a TOC90G-SEP1G heterodimer and TOC90G-TOC34G heterodimer. (F) Cartoon schematic of FragFold workflow used in panel E. (G-H) Predicted TOC90-SEP1G (G) and TOC90G-TOC34G (H) contact sites on TOC90G monomer. GTP-binding face is indicated. Color scaled by mean weighted contact score.

**Supplementary Figure 9.**
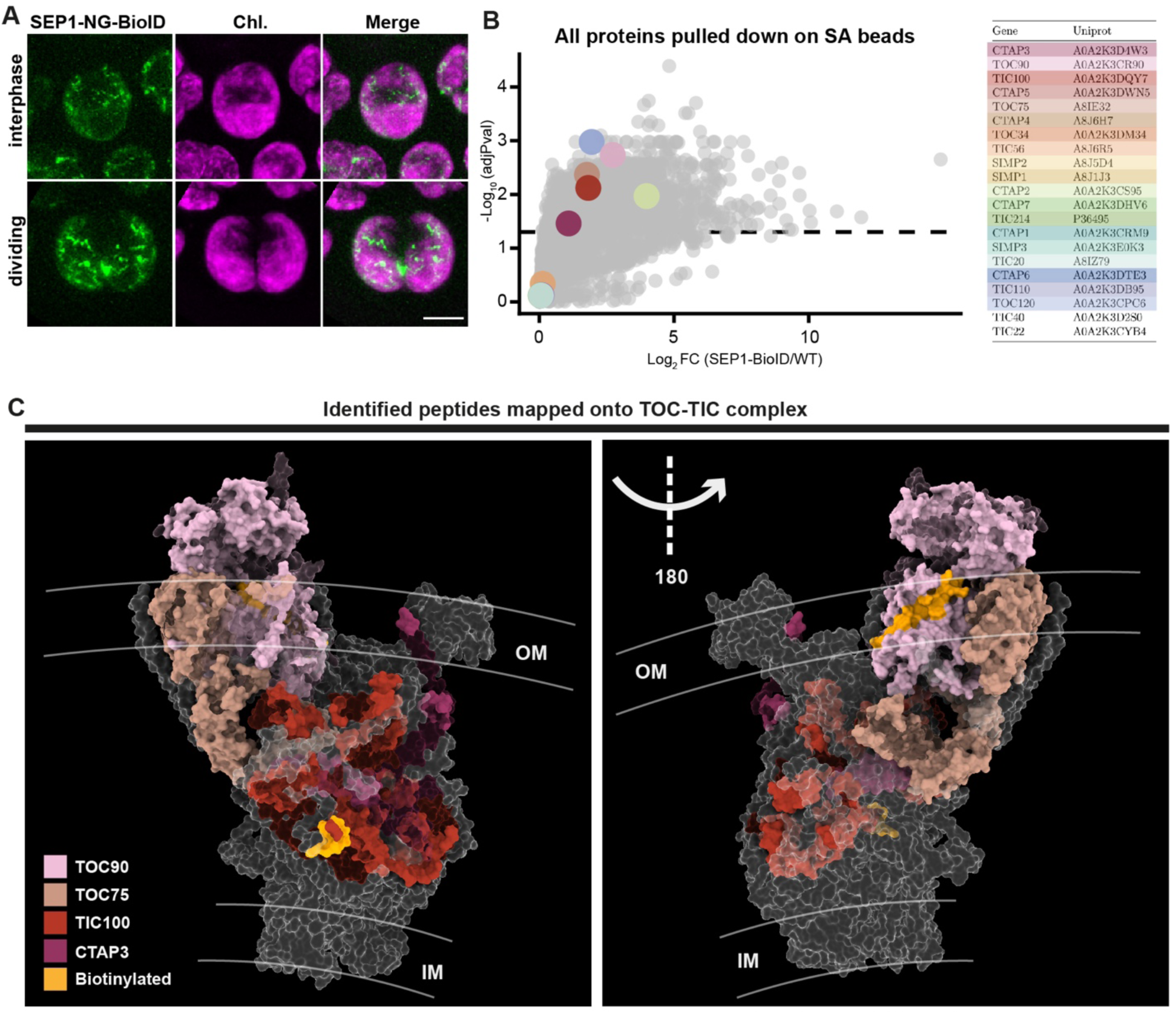
SEP1-NG-BioIDG3 localizes to the chloroplast surface and identifies TOC proteins. (A) Live cell microscopy of SEP1-NG-BioIDG3 strain in interphase and dividing cells. Maximum Z projections. Scale bar: 5 μm. (B) Volcano plot of identified chloroplast translocon components from the proximity-based labeling experiment. Dotted line marks FDR-adjusted P = 0.05 (LIMMA empirical Bayes-moderated t-test with Benjamini-Hochberg correction). (C) Positions of proteins identified by SEP1-BioIDG3 on the TOC-TIC structure (PDB: 7VCF). Colors of subunits match those in (B). Because only the transmembrane domain of TOC90 was resolved in the structure, an AlphaFold-predicted structure of full-length TOC90 (light pink) is superimposed. Yellow regions represent identified biotinylated peptides.

**Supplementary Figure 10.**
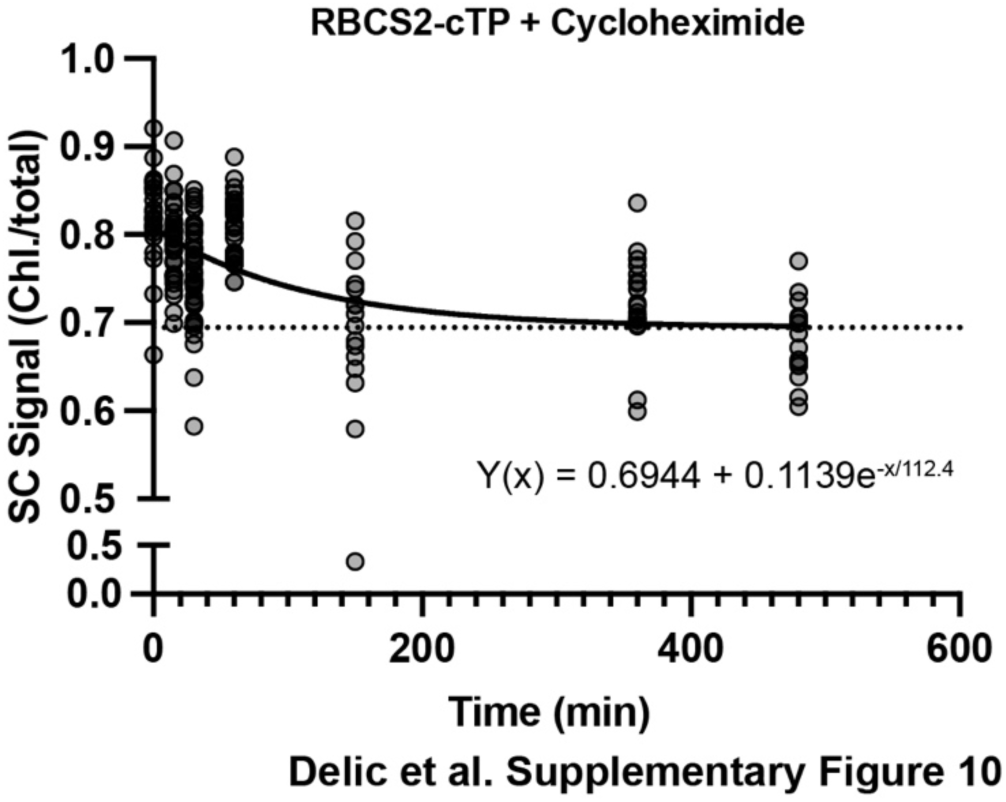
Cycloheximide treatment establishes baseline for cTP reporter assays. 30 µg/mL cycloheximide was added at 0 min to an exponentially growing culture of cells expressing cTP^RBCS2^-SC reporter. Fluorescence intensity measurements were taken at indicated time points, and ratiometric quantification was done as outlined in Figure 4C. Dotted line corresponds to the baseline ratio from autofluorescence.

**Supplementary Figure 11.**
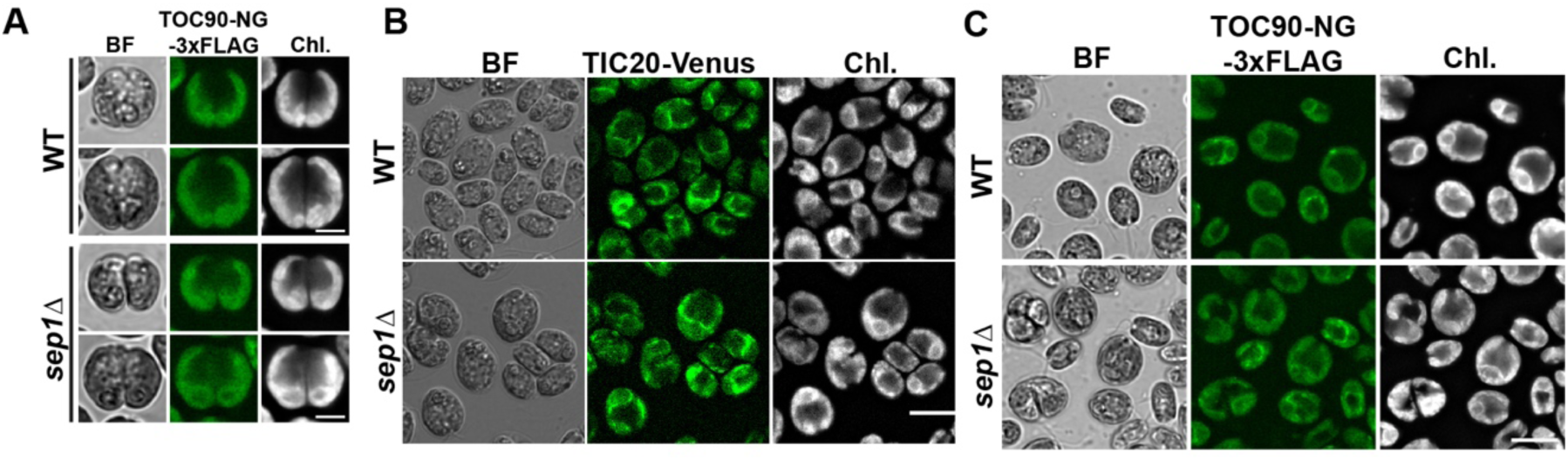
Loss of SEP1 does not alter translocon localization or apparent abundance. (A) Localization of TOC90-NG-3FLAG in dividing WT and *sep1Δ* cells. Maximum Z projection. Scale bar: 5 μm. (B) Localization of TIC20-Venus in WT and *sep1Δ* cells. Single Z slice. Scale bar: 10 μm. (C) Larger field of view of TOC90-NG-3FLAG in WT and *sep1Δ* cells. Average Z projection. Scale bar: 5 μm.

**Supplementary Figure 12.**
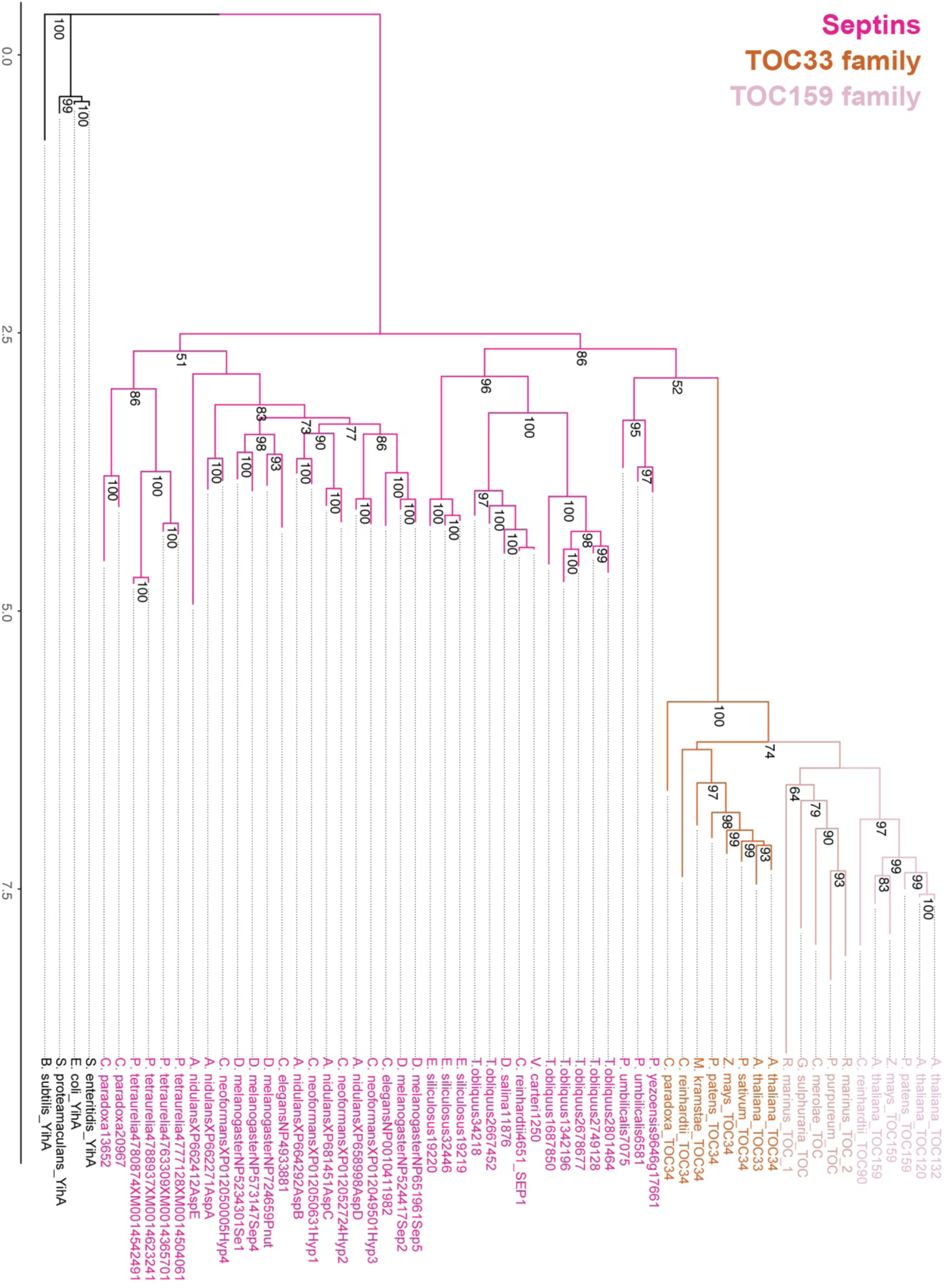
Phylogenetic tree of septins and TOC GTPases shown in. **Figure 5A**. Maximum likelihood tree of 39 septin sequences with 20 TOC GTPases, including putative TOC GTPases from early archeaplastida lineages. Branch lengths displayed on x-axis. Bootstrap values (0-100) displayed at nodes. Individual septin, and TOC GTPase families are color coded. Tree is rooted using 4 prokaryotic YihA proteins.

**Supplementary Figure 13.**
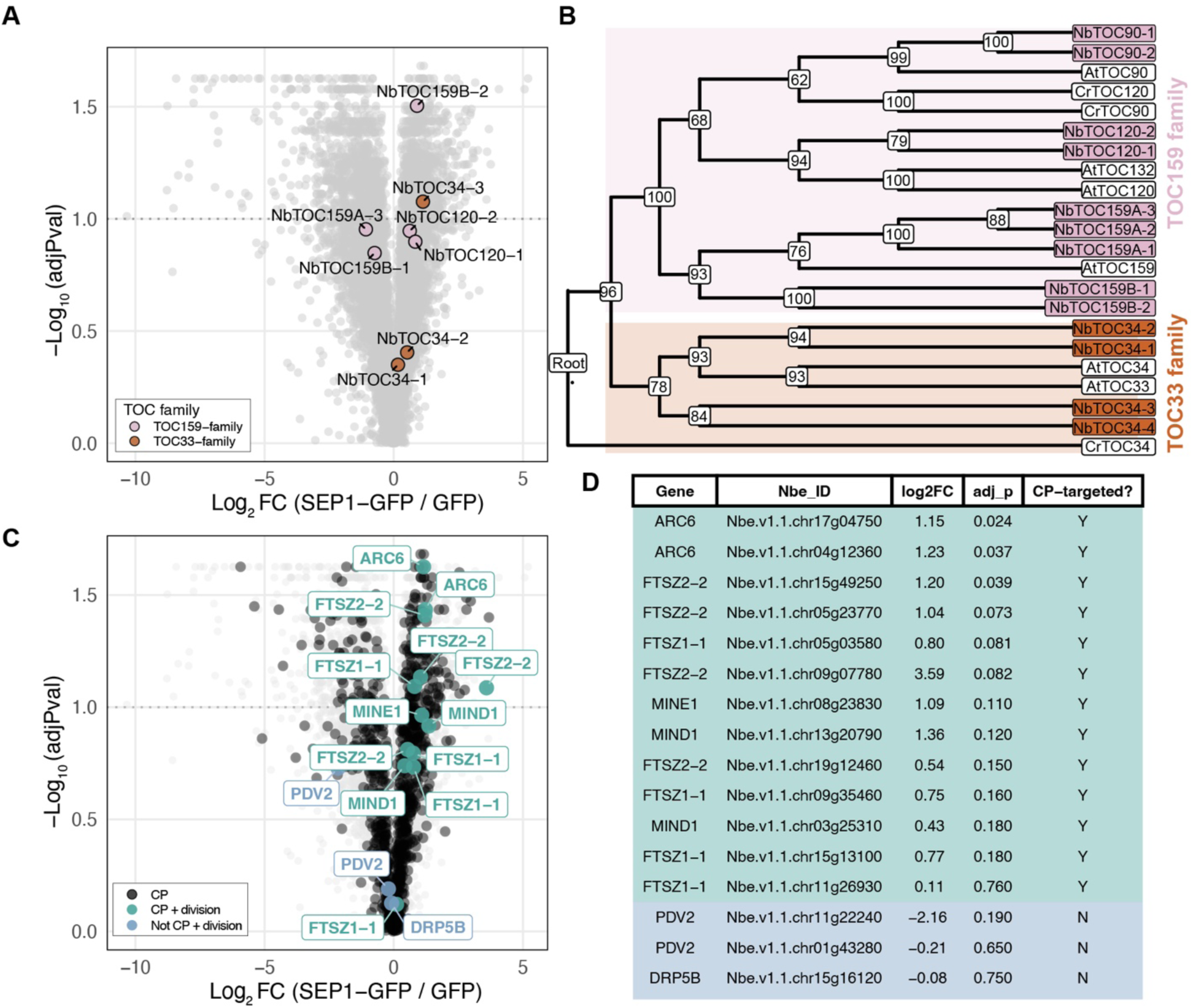
SEP1 binds to chloroplast translocons in land plants. (A) Volcano plot of mass spectrometry hits from *N. benthamiana* for CrSEP1-GFP immunoprecipitation compared to GFP-alone control. Dotted line represents the FDR-adjusted P = 0.1 (t-test with Benjamini-Hochberg correction). Colored points represent identified TOC159-family and TOC33-family member proteins. (B) Cladogram of identified TOC GTPases in the *N. benthamiana* genome. (C) Volcano plot of mass spectrometry hits highlighting chloroplast-targeted and chloroplast-division proteins. (D) Table of identified chloroplast division-proteins identified in (C). Chloroplast-targeted division proteins are shown in green. Non-chloroplast-targeted division proteins are shown in blue.

## REFERENCES

1. Sagan, L. (1967). On the origin of mitosing cells. Journal of Theoretical Biology 14, 225-IN6. 10.1016/0022-5193(67)90079-3.

2. Martin, W., and Kowallik, K.V. (1999). Annotated English translation of Mereschkowsky’s 1905 paper ‘Über Natur und Ursprung der Chromatophoren im Pflanzenreiche.’ European Journal of Phycology 34, 287–295.

3. Cavalier-Smith, T. (1987). The simultaneous symbiotic origin of mitochondria, chloroplasts, and microbodies. Annals of the New York Academy of Sciences 503, 55–71. 10.1111/j.1749-6632.1987.tb40597.x.

4. Lyons, T.W., Reinhard, C.T., and Planavsky, N.J. (2014). The rise of oxygen in Earth’s early ocean and atmosphere. Nature 506, 307–315. 10.1038/nature13068.

5. Sánchez-Baracaldo, P., Bianchini, G., Wilson, J.D., and Knoll, A.H. (2022). Cyanobacteria and biogeochemical cycles through Earth history. Trends in Microbiology 30, 143–157. 10.1016/j.tim.2021.05.008.

6. Sumiya, N., Fujiwara, T., Era, A., and Miyagishima, S. (2016). Chloroplast division checkpoint in eukaryotic algae. Proceedings of the National Academy of Sciences 113, E7629–E7638. 10.1073/pnas.1612872113.

7. de Vries, J., and Gould, S.B. (2018). The monoplastidic bottleneck in algae and plant evolution. J Cell Sci 131, jcs203414. 10.1242/jcs.203414.

8. Martin, W., Rujan, T., Richly, E., Hansen, A., Cornelsen, S., Lins, T., Leister, D., Stoebe, B., Hasegawa, M., and Penny, D. (2002). Evolutionary analysis of *Arabidopsis*, cyanobacterial, and chloroplast genomes reveals plastid phylogeny and thousands of cyanobacterial genes in the nucleus. Proceedings of the National Academy of Sciences 99, 12246–12251. 10.1073/pnas.182432999.

9. Martin, W., Stoebe, B., Goremykin, V., Hansmann, S., Hasegawa, M., and Kowallik, K.V. (1998). Gene transfer to the nucleus and the evolution of chloroplasts. Nature 393, 162–165. 10.1038/30234.

10. Schnell, D.J., Kessler, F., and Blobel, G. (1994). Isolation of components of the chloroplast protein import machinery. Science 266, 1007–1012. 10.1126/science.7973649.

11. Bauer, J., Chen, K., Hiltbunner, A., Wehrli, E., Eugster, M., Schnell, D., and Kessler, F. (2000). The major protein import receptor of plastids is essential for chloroplast biogenesis. Nature 403, 203–207. 10.1038/35003214.

12. Jarvis, P., Chen, L.-J., Li, H., Peto, C.A., Fankhauser, C., and Chory, J. (1998). An *Arabidopsis* mutant defective in the plastid general protein import apparatus. Science 282, 100–103. 10.1126/science.282.5386.100.

13. Rodríguez-Ezpeleta, N., and Philippe, H. (2006). Plastid origin: replaying the tape. Current Biology 16, R53–R56. 10.1016/j.cub.2006.01.006.

14. Okazaki, K., Kabeya, Y., and Miyagishima, S. (2010). The evolution of the regulatory mechanism of chloroplast division. Plant Signal Behav 5, 164–167.

15. Yoshida, Y., Mogi, Y., TerBush, A.D., and Osteryoung, K.W. (2016). Chloroplast FtsZ assembles into a contractible ring via tubulin-like heteropolymerization. Nat Plants 2, 16095. 10.1038/nplants.2016.95.

16. Gremillon, L., Kiessling, J., Hause, B., Decker, E.L., Reski, R., and Sarnighausen, E. (2007). Filamentous temperature-sensitive Z (FtsZ) isoforms specifically interact in the chloroplasts and in the cytosol of *Physcomitrella patens*. New Phytol 176, 299–310. 10.1111/j.1469-8137.2007.02169.x.

17. Wang, D., Kong, D., Wang, Y., Hu, Y., He, Y., and Sun, J. (2003). Isolation of two plastid division ftsZ genes from *Chlamydomonas reinhardtii* and its evolutionary implication for the role of FtsZ in plastid division. J Exp Bot 54, 1115–1116. 10.1093/jxb/erg117.

18. Irieda, H., and Shiomi, D. (2017). ARC6-mediated Z ring-like structure formation of prokaryote-descended chloroplast FtsZ in *Escherichia coli*. Sci Rep 7, 3492. 10.1038/s41598-017-03698-6.

19. 19. Vitha, S., Froehlich, J.E., Koksharova, O., Pyke, K.A., van Erp, H., and Osteryoung, K.W. (2003). ARC6 is a J-domain plastid division protein and an evolutionary descendant of the cyanobacterial cell division protein Ftn2. Plant Cell 15, 1918–1933. 10.1105/tpc.013292.

20. Johnson, C.B., Tang, L.K., Smith, A.G., Ravichandran, A., Luo, Z., Vitha, S., and Holzenburg, A. (2013). Single particle tracking analysis of the chloroplast division protein FtsZ anchoring to the inner envelope membrane. Microscopy and Microanalysis 19, 507–512. 10.1017/S143192761300038X.

21. Osteryoung, K.W., and Vierling, E. (1995). Conserved cell and organelle division. Nature 376, 473–474. 10.1038/376473b0.

22. Fujiwara, M., and Yoshida, S. (2001). Chloroplast targeting of chloroplast division FtsZ2 proteins in *Arabidopsis*. Biochem Biophys Res Commun 287, 462–467. 10.1006/bbrc.2001.5588.

23. Keegstra, K. (1989). Transport and routing of proteins into chloroplasts. Cell 56, 247–253. 10.1016/0092-8674(89)90898-2.

24. Mishkind, M.L., Wessler, S.R., and Schmidt, G.W. (1985). Functional determinants in transit sequences: import and partial maturation by vascular plant chloroplasts of the ribulose-1,5-bisphosphate carboxylase small subunit of *Chlamydomonas*. Journal of Cell Biology 100, 226–234. 10.1083/jcb.100.1.226.

25. Nellaepalli, S., Baretic, D., Brandner, A.F., Kubis-Waller, S., Soufi, Z., Cheng, S., Kodru, S., Fang, J., Flores-Perez, U., Ravikumar, V., et al. (2025). Assembly of two functionally-distinct protein import complexes in the outer membrane of plant chloroplasts. Preprint at bioRxiv, 10.1101/2025.10.27.683105

26. Constan, D., Patel, R., Keegstra, K., and Jarvis, P. (2004). An outer envelope membrane component of the plastid protein import apparatus plays an essential role in *Arabidopsis*. The Plant Journal 38, 93–106. 10.1111/j.1365-313X.2004.02024.x.

27. Reumann, S., Davila-Aponte, J., and Keegstra, K. (1999). The evolutionary origin of the protein-translocating channel of chloroplastic envelope membranes: Identification of a cyanobacterial homolog. Proceedings of the National Academy of Sciences 96, 784–789. 10.1073/pnas.96.2.784.

28. Reumann, S., Inoue, K., and Keegstra, K. (2005). Evolution of the general protein import pathway of plastids (Review). Molecular Membrane Biology 22, 73–86. 10.1080/09687860500041916.

29. Shanmugabalaji, V., Chahtane, H., Accossato, S., Rahire, M., Gouzerh, G., Lopez-Molina, L., and Kessler, F. (2018). Chloroplast biogenesis controlled by DELLA-TOC159 interaction in early plant development. Curr Biol 28, 2616–2623.e5. 10.1016/j.cub.2018.06.006.

30. Fang, J., Li, B., Chen, L.-J., Dogra, V., Luo, S., Wu, W., Wang, P., Hwang, I., Li, H., and Kim, C. (2022). TIC236 gain-of-function mutations unveil the link between plastid division and plastid protein import. Proceedings of the National Academy of Sciences 119, e2123353119. 10.1073/pnas.2123353119.

31. Zhang, J., Wu, S., Boehlein, S.K., McCarty, D.R., Song, G., Walley, J.W., Myers, A., and Settles, A.M. (2019). Maize defective kernel5 is a bacterial TamB homologue required for chloroplast envelope biogenesis. J Cell Biol 218, 2638–2658. 10.1083/jcb.201807166.

32. Field, C.M., al-Awar, O., Rosenblatt, J., Wong, M.L., Alberts, B., and Mitchison, T.J. (1996). A purified *Drosophila* septin complex forms filaments and exhibits GTPase activity. The Journal of cell biology 133, 605–616. 10.1083/jcb.133.3.605.

33. Sirajuddin, M., Farkasovsky, M., Hauer, F., Kühlmann, D., Macara, I.G., Weyand, M., Stark, H., and Wittinghofer, A. (2007). Structural insight into filament formation by mammalian septins. Nature 449, 311–315. 10.1038/nature06052.

34. Byers, B., and Goetsch, L. (1976). A highly ordered ring of membrane-associated filaments in budding yeast. Journal of Cell Biology 69, 717–721. 10.1083/jcb.69.3.717.

35. Bridges, A.A., Jentzsch, M.S., Oakes, P.W., Occhipinti, P., and Gladfelter, A.S. (2016). Micron-scale plasma membrane curvature is recognized by the septin cytoskeleton. J Cell Biol 213, 23–32. 10.1083/jcb.201512029.

36. McMurray, M.A., Bertin, A., Garcia, G., Lam, L., Nogales, E., and Thorner, J. (2011). Septin Filament Formation is Essential in Budding Yeast. Dev Cell 20, 540–549. 10.1016/j.devcel.2011.02.004.

37. Bertin, A., McMurray, M.A., Thai, L., Garcia, G., Votin, V., Grob, P., Allyn, T., Thorner, J., and Nogales, E. (2010). Phosphatidylinositol-4,5-bisphosphate promotes budding yeast septin filament assembly and organization. Journal of Molecular Biology 404, 711–731. 10.1016/j.jmb.2010.10.002.

38. Pagliuso, A., Tham, T.N., Stevens, J.K., Lagache, T., Persson, R., Salles, A., Olivo-Marin, J.-C., Oddos, S., Spang, A., Cossart, P., et al. (2016). A role for septin 2 in Drp1-mediated mitochondrial fission. EMBO Rep 17, 858–873. 10.15252/embr.201541612.

39. Sirianni, A., Krokowski, S., Lobato-Márquez, D., Buranyi, S., Pfanzelter, J., Galea, D., Willis, A., Culley, S., Henriques, R., Larrouy-Maumus, G., et al. (2016). Mitochondria mediate septin cage assembly to promote autophagy of *Shigella*. EMBO Rep 17, 1029–1043. 10.15252/embr.201541832.

40. Mageswaran, S.K., Grotjahn, D.A., Zeng, X., Barad, B.A., Medina, M., Hoang, M.H., Dobro, M.J., Chang, Y.-W., Xu, M., Yang, W.Y., et al. (2023). Nanoscale details of mitochondrial constriction revealed by cryoelectron tomography. Biophysical Journal 122, 3768–3782. 10.1016/j.bpj.2023.07.030.

41. Huang, Y.-W., Yan, M., Collins, R.F., DiCiccio, J.E., Grinstein, S., and Trimble, W.S. (2008). Mammalian septins are required for phagosome formation. Mol Biol Cell 19, 1717–1726. 10.1091/mbc.E07-07-0641.

42. Mostowy, S., Bonazzi, M., Hamon, M.A., Tham, T.N., Mallet, A., Lelek, M., Gouin, E., Demangel, C., Brosch, R., Zimmer, C., et al. (2010). Entrapment of intracytosolic bacteria by septin cage-like structures. Cell Host & Microbe 8, 433–444. 10.1016/j.chom.2010.10.009.

43. Yamazaki, T., Owari, S., Ota, S., Sumiya, N., Yamamoto, M., Watanabe, K., Nagumo, T., Miyamura, S., and Kawano, S. (2013). Localization and evolution of septins in algae. The Plant Journal 74, 605–614. 10.1111/tpj.12147.

44. Wloga, D., Strzyzewska-Jówko, I., Gaertig, J., and Jerka-Dziadosz, M. (2008). Septins stabilize mitochondria in *Tetrahymena thermophila*. Eukaryot Cell 7, 1373–1386. 10.1128/EC.00085-08.

45. Pinto, A.P.A., Pereira, H.M., Zeraik, A.E., Ciol, H., Ferreira, F.M., Brandão-Neto, J., DeMarco, R., Navarro, M.V.A.S., Risi, C., Galkin, V.E., et al. (2017). Filaments and fingers: novel structural aspects of the single septin from *Chlamydomonas reinh*ardtii. J Biol Chem 292, 10899–10911. 10.1074/jbc.M116.762229.

46. Onishi, M., and Pringle, J.R. (2016). The nonopisthokont septins: how many there are, how little we know about them, and how we might learn more. Methods Cell Biol 136, 1–19. 10.1016/bs.mcb.2016.04.003.

47. Goodson, H.V., Kelley, J.B., and Brawley, S.H. (2021). Cytoskeletal diversification across 1 billion years: what red algae can teach us about the cytoskeleton, and vice versa. BioEssays 43, 2000278. 10.1002/bies.202000278.

48. Delic, S., Shuman, B., Lee, S., Bahmanyar, S., Momany, M., and Onishi, M. (2024). The evolutionary origins and ancestral features of septins. Front. Cell Dev. Biol. 12. 10.3389/fcell.2024.1406966.

49. Versele, M., and Thorner, J. (2005). Some assembly required: yeast septins provide the instruction manual. Trends Cell Biol 15, 414–424. 10.1016/j.tcb.2005.06.007.

50. Cannon, K.S., Woods, B.L., Crutchley, J.M., and Gladfelter, A.S. (2019). An amphipathic helix enables septins to sense micrometer-scale membrane curvature. Journal of Cell Biology 218, 1128–1137. 10.1083/jcb.201807211.

51. Cross, F.R., and Umen, J.G. (2015). The *Chlamydomonas* cell cycle. The Plant Journal 82, 370–392. 10.1111/tpj.12795.

52. Gaffal, K.P., Arnold, C.-G., Friedrichs, G.J., and Gemple, W. (1995). Morphodynamical changes of the chloroplast of *Chlamydomonas reinhardtii* during the 1st round of division. Archiv für Protistenkunde 145, 10–23. 10.1016/S0003-9365(11)80297-6.

53. 53. Nakazawa, K., Kumar, G., Chauvin, B., Di Cicco, A., Pellegrino, L., Trichet, M., Hajj, B., Cabral, J., Sain, A., Mangenot, S., et al. (2023). A human septin octamer complex sensitive to membrane curvature drives membrane deformation with a specific mesh-like organization. J Cell Sci 136, jcs260813. 10.1242/jcs.260813.

54. Woods, B.L., Cannon, K.S., Vogt, E.J.D., Crutchley, J.M., and Gladfelter, A.S. (2021). Interplay of septin amphipathic helices in sensing membrane-curvature and filament bundling. Mol Biol Cell 32, br5. 10.1091/mbc.E20-05-0303.

55. Onishi, M., Umen, J.G., Cross, F.R., and Pringle, J.R. (2020). Cleavage-furrow formation without F-actin in *Chlamydomonas*. Proceedings of the National Academy of Sciences 117, 18511–18520. 10.1073/pnas.1920337117.

56. Clark-Cotton, M.R., Chen, S.-A., Gomez, A., Mulabagal, A.J., Perry, A., Malhotra, V., and Onishi, M. (2025). Imaging-based screen identifies novel natural compounds that perturb cell and chloroplast division in *Chlamydomonas reinhardtii*. MBoC 36, br14. 10.1091/mbc.E24-09-0425.

57. Onishi, M., Pecani, K., Jones, T., Pringle, J.R., and Cross, F.R. (2018). F-actin homeostasis through transcriptional regulation and proteasome-mediated proteolysis. Proceedings of the National Academy of Sciences 115, E6487–E6496. 10.1073/pnas.1721935115.

58. Onishi, M., Pringle, J.R., and Cross, F.R. (2016). Evidence that an unconventional actin can provide essential F-actin function and that a surveillance system monitors F-actin integrity in *Chlamydomonas*. Genetics 202, 977–996. 10.1534/genetics.115.184663.

59. Avasthi, P., and Onishi, M. (2023). The cellular cytoskeleton. In The Chlamydomonas Sourcebook (Elsevier), pp. 433–445. 10.1016/B978-0-12-822508-0.00001-0.

60. Liu, H., Li, A., Rochaix, J.-D., and Liu, Z. (2023). Architecture of chloroplast TOC–TIC translocon supercomplex. Nature 615, 349–357. 10.1038/s41586-023-05744-y.

61. Jin, Z., Wan, L., Zhang, Y., Li, X., Cao, Y., Liu, H., Fan, S., Cao, D., Wang, Z., Li, X., et al. (2022). Structure of a TOC-TIC supercomplex spanning two chloroplast envelope membranes. Cell 185, 4788–4800.e13. 10.1016/j.cell.2022.10.030.

62. Wang, N., Xing, J., Su, X., Pan, J., Chen, H., Shi, L., Si, L., Yang, W., and Li, M. (2024). Architecture of the ATP-driven motor for protein import into chloroplasts. Molecular Plant 17, 1702–1718. 10.1016/j.molp.2024.09.010.

63. Weibel, P., Hiltbrunner, A., Brand, L., and Kessler, F. (2003). Dimerization of Toc-GTPases at the chloroplast protein import machinery. Journal of Biological Chemistry 278, 37321–37329. 10.1074/jbc.M305946200.

64. Becker, T., Jelic, M., Vojta, A., Radunz, A., Soll, J., and Schleiff, E. (2004). Preprotein recognition by the Toc complex. EMBO J 23, 520–530. 10.1038/sj.emboj.7600089.

65. Richter, S., and Lamppa, G.K. (1999). Stromal processing peptidase binds transit peptides and initiates their ATP-dependent turnover in chloroplasts. J Cell Biol 147, 33–44.

66. Weirich, C.S., Erzberger, J.P., and Barral, Y. (2008). The septin family of GTPases: architecture and dynamics. Nat Rev Mol Cell Biol 9, 478–489. 10.1038/nrm2407.

67. Ramundo, S., Asakura, Y., Salomé, P.A., Strenkert, D., Boone, M., Mackinder, L.C.M., Takafuji, K., Dinc, E., Rahire, M., Crèvecoeur, M., et al. (2020). Coexpressed subunits of dual genetic origin define a conserved supercomplex mediating essential protein import into chloroplasts. Proceedings of the National Academy of Sciences 117, 32739–32749. 10.1073/pnas.2014294117.

68. Zones, J.M., Blaby, I.K., Merchant, S.S., and Umen, J.G. (2015). High-resolution profiling of a synchronized diurnal transcriptome from *Chlamydomonas reinhardtii* reveals continuous cell and metabolic differentiation. Plant Cell 27, 2743–2769. 10.1105/tpc.15.00498.

69. Leipe, D.D., Wolf, Y.I., Koonin, E.V., and Aravind, L. (2002). Classification and evolution of P-loop GTPases and related ATPases. J Mol Biol 317, 41–72. 10.1006/jmbi.2001.5378.

70. Engel, B.D., Schaffer, M., Kuhn Cuellar, L., Villa, E., Plitzko, J.M., and Baumeister, W. (2015). Native architecture of the *Chlamydomonas* chloroplast revealed by in situ cryo-electron tomography. eLife 4, e04889. 10.7554/eLife.04889.

71. Liang, K., Zhan, X., Li, Y., Yang, Y., Xie, Y., Jin, Z., Xu, X., Zhang, W., Lu, Y., Zhang, S., et al. (2024). Conservation and specialization of the Ycf2-FtsHi chloroplast protein import motor in green algae. Cell 187, 5638–5650.e18. 10.1016/j.cell.2024.08.002.

72. Kim, M.S., Froese, C.D., Xie, H., and Trimble, W.S. (2012). Uncovering principles that control septin-septin interactions. J Biol Chem 287, 30406–30413. 10.1074/jbc.M112.387464.

73. Kinoshita, M., Field, C.M., Coughlin, M.L., Straight, A.F., and Mitchison, T.J. (2002). Self-and actin-templated assembly of mammalian septins. Developmental Cell 3, 791–802. 10.1016/S1534-5807(02)00366-0.

74. 74. Estey, M.P., Di Ciano-Oliveira, C., Froese, C.D., Bejide, M.T., and Trimble, W.S. (2010). Distinct roles of septins in cytokinesis: SEPT9 mediates midbody abscission. J Cell Biol 191, 741–749. 10.1083/jcb.201006031.

75. Gonzalez, M.E., Makarova, O., Peterson, E.A., Privette, L.M., and Petty, E.M. (2009). Up-regulation of SEPT9_v1 stabilizes c-Jun-N-Terminal kinase and contributes to its pro-proliferative activity in mammary epithelial cells. Cellular Signalling 21, 477–487. 10.1016/j.cellsig.2008.11.007.

76. Oreb, M., Höfle, A., Koenig, P., Sommer, M.S., Sinning, I., Wang, F., Tews, I., Schnell, D.J., and Schleiff, E. (2011). Substrate binding disrupts dimerization and induces nucleotide exchange of the chloroplast GTPase Toc33. Biochem J 436, 313–319. 10.1042/BJ20110246.

77. Kubis, S., Baldwin, A., Patel, R., Razzaq, A., Dupree, P., Lilley, K., Kurth, J., Leister, D., and Jarvis, P. (2003). The *Arabidopsis* ppi1 mutant is specifically defective in the expression, chloroplast import, and accumulation of photosynthetic proteins. Plant Cell 15, 1859–1871. 10.1105/tpc.012955.

78. Ford, S.K., and Pringle, J.R. (1991). Cellular morphogenesis in the *Saccharomyces cerevisiae* cell cycle: localization of the CDC11 gene product and the timing of events at the budding site. Dev Genet 12, 281–292. 10.1002/dvg.1020120405.

79. Haarer, B.K., and Pringle, J.R. (1987). Immunofluorescence localization of the *Saccharomyces cerevisiae* CDC12 gene product to the vicinity of the 10-nm filaments in the mother-bud neck. Mol Cell Biol 7, 3678–3687. 10.1128/mcb.7.10.3678-3687.1987.

80. Hartwell, L.H. (1971). Genetic control of the cell division cycle in yeast: IV. Genes controlling bud emergence and cytokinesis. Experimental Cell Research 69, 265–276. 10.1016/0014-4827(71)90223-0.

81. Neufeld, T.P., and Rubin, G.M. (1994). The *Drosophila* peanut gene is required for cytokinesis and encodes a protein similar to yeast putative bud neck filament proteins. Cell 77, 371–379. 10.1016/0092-8674(94)90152-X.

82. Kinoshita, M., Kumar, S., Mizoguchi, A., Ide, C., Kinoshita, A., Haraguchi, T., Hiraoka, Y., and Noda, M. (1997). Nedd5, a mammalian septin, is a novel cytoskeletal component interacting with actin-based structures. Genes Dev. 11, 1535–1547. 10.1101/gad.11.12.1535.

83. Nguyen, T.Q., Sawa, H., Okano, H., and White, J.G. (2000). The *C. elegans* septin genes, unc-59 and unc-61, are required for normal postembryonic cytokineses and morphogenesis but have no essential function in embryogenesis. J Cell Sci 113, 3825–3837. 10.1242/jcs.113.21.3825.

84. Pyke, K.A., and Leech, R.M. (1994). A genetic analysis of chloroplast division and expansion in *Arabidopsis thaliana*. Plant Physiol 104, 201–207. 10.1104/pp.104.1.201.

85. Bai, X., Bowen, J.R., Knox, T.K., Zhou, K., Pendziwiat, M., Kuhlenbäumer, G., Sindelar, C.V., and Spiliotis, E.T. (2013). Novel septin 9 repeat motifs altered in neuralgic amyotrophy bind and bundle microtubules. J Cell Biol 203, 895–905. 10.1083/jcb.201308068.

86. Mavrakis, M., Azou-Gros, Y., Tsai, F.-C., Alvarado, J., Bertin, A., Iv, F., Kress, A., Brasselet, S., Koenderink, G.H., and Lecuit, T. (2014). Septins promote F-actin ring formation by crosslinking actin filaments into curved bundles. Nat Cell Biol 16, 322–334. 10.1038/ncb2921.

87. Nakos, K., Alam, M.N.A., Radler, M.R., Kesisova, I.A., Yang, C., Okletey, J., Tomasso, M.R., Padrick, S.B., Svitkina, T.M., and Spiliotis, E.T. (2022). Septins mediate a microtubule–actin crosstalk that enables actin growth on microtubules. Proceedings of the National Academy of Sciences 119, e2202803119. 10.1073/pnas.2202803119.

88. Oh, Y., and Bi, E. (2011). Septin structure and function in yeast and beyond. Trends Cell Biol 21, 141–148. 10.1016/j.tcb.2010.11.006.

89. Shannon, R., Balachandran, Y., Wang, X., Boutry, M., Xie, H., Kim, P.K., and Trimble, W.S. (2025). SEPTIN9 locally activates the RhoGEF ARHGEF18 to promote early stages of mitochondrial fission. J Cell Biol 224, e202406017. 10.1083/jcb.202406017.

90. Perry, J.A., Werner, M.E., Omi, S., Heck, B.W., Maddox, P.S., Mavrakis, M., and Maddox, A.S. (2025). Animal septins contain functional transmembrane domains. Current Biology 35, 1910–1917.e5. 10.1016/j.cub.2025.03.004.

91. 91. Dutcher, S.K. (1995). Chapter 76 Mating and Tetrad Analysis in Chlamydomonas reinhardtii. In Methods in Cell Biology (Academic Press), pp. 531–540. 10.1016/S0091-679X(08)60857-2.

92. Onishi, M., and Pringle, J.R. (2016). Robust transgene expression from bicistronic mRNA in the green alga *Chlamydomonas reinhardtii*. G3 (Bethesda) 6, 4115–4125. 10.1534/g3.116.033035.

93. Wang, L., Patena, W., Baalen, K.A.V., Xie, Y., Singer, E.R., Gavrilenko, S., Warren-Williams, M., Han, L., Harrigan, H.R., Hartz, L.D., et al. (2023). A chloroplast protein atlas reveals punctate structures and spatial organization of biosynthetic pathways. Cell 186, 3499–3518.e14. 10.1016/j.cell.2023.06.008.

94. Laporte, M.H., Klena, N., Hamel, V., and Guichard, P. (2022). Visualizing the native cellular organization by coupling cryofixation with expansion microscopy (Cryo-ExM). Nat Methods 19, 216–222. 10.1038/s41592-021-01356-4.

95. 95. Gambarotto, D., Zwettler, F.U., Le Guennec, M., Schmidt-Cernohorska, M., Fortun, D., Borgers, S., Heine, J., Schloetel, J.-G., Reuss, M., Unser, M., et al. (2019). Imaging cellular ultrastructures using expansion microscopy (U-ExM). Nat Methods 16, 71–74. 10.1038/s41592-018-0238-1.

96. Fields, S., and Song, O. (1989). A novel genetic system to detect protein-protein interactions. Nature 340, 245–246. 10.1038/340245a0.

97. Cox, J., Hein, M.Y., Luber, C.A., Paron, I., Nagaraj, N., and Mann, M. (2014). Accurate Proteome-wide label-free quantification by delayed normalization and maximal peptide ratio extraction, Termed MaxLFQ. Molecular & Cellular Proteomics 13, 2513–2526. 10.1074/mcp.M113.031591.

98. Huang, L.-K., and Wang, M.-J.J. (1995). Image thresholding by minimizing the measures of fuzziness. Pattern Recognition 28, 41–51. 10.1016/0031-3203(94)E0043-K.

99. 99. Starr, R.C. (1969). Structure, reproduction and differentiation in *Volvox carteri f. nagariensis* Iyengar, Strains HK 9 & 10. In Archiv für Protistenkunde, pp. 204–222.

100. 100. Tian, Y., Gao, S., von der Heyde, E.L., Hallmann, A., and Nagel, G. (2018). Two-component cyclase opsins of green algae are ATP-dependent and light-inhibited guanylyl cyclases. BMC Biol 16, 144. 10.1186/s12915-018-0613-5.

101. 101. Provasoli, L., Pintner, I. J. (1959) Artificial media for fresh-water algae: problems and suggestions. In: Tryon, C. A. und Hartman, R. T. (Hg.): The Ecology of Algae. a symposium held at the Pymatuning Laboratory of Field Biology. Pittsburgh, PA, June 18 and 19, 1959 (The Pymatuning Symposia, Special Publication No. 2), S. 84–96. University of Pittsburgh. 10.5962/bhl.title.5728

102. 102. von der Heyde, B., and Hallmann, A. (2022). Cell type-specific pherophorins of *Volvox carteri* reveal interplay of both cell types in ECM biosynthesis. Cells 12, 134. 10.3390/cells12010134.

103. Starr, R.C., and Jaenicke, L. (1974). Purification and characterization of the hormone initiating sexual morphogenesis in *Volvox carteri f. nagariensis* Iyengar. Proceedings of the National Academy of Sciences 71, 1050–1054. 10.1073/pnas.71.4.1050.

104. 104. von der Heyde, E.L., and Hallmann, A. (2022). Molecular and cellular dynamics of early embryonic cell divisions in Volvox carteri. Plant Cell 34, 1326–1353. 10.1093/plcell/koac004.

105. Hallmann, A., and Wodniok, S. (2006). Swapped green algal promoters: aphVIII-based gene constructs with *Chlamydomonas* flanking sequences work as dominant selectable markers in *Volvox* and vice versa. Plant Cell Rep 25, 582–591. 10.1007/s00299-006-0121-x.

106. Schiedlmeier, B., Schmitt, R., Müller, W., Kirk, M.M., Gruber, H., Mages, W., and Kirk, D.L. (1994). Nuclear transformation of *Volvox carteri*. Proceedings of the National Academy of Sciences 91, 5080–5084. 10.1073/pnas.91.11.5080.

107. Gruber, H., Kirzinger, S.H., and Schmitt, R. (1996). Expression of the *Volvox* gene encoding nitrate reductase: mutation-dependent activation of cryptic splice sites and intron-enhanced gene expression from a cDNA. Plant Mol Biol 31, 1–12. 10.1007/BF00020601.

108. 108. von der Heyde, E.L., Klein, B., Abram, L., and Hallmann, A. (2015). The inducible nitA promoter provides a powerful molecular switch for transgene expression in *Volvox carteri*. BMC Biotechnol 15, 5. 10.1186/s12896-015-0122-3.

109. 109. von der Heyde, B., Srinivasan, A., Birwa, S.K., von der Heyde, E.L., Höhn, S.S.M.H., Goldstein, R.E., and Hallmann, A. (2025). Spatiotemporal distribution of the glycoprotein pherophorin II reveals stochastic geometry of the growing ECM of *Volvox carteri*. Proc Natl Acad Sci U S A 122, e2425759122. 10.1073/pnas.2425759122.

110. Wu, S.-Z., Ryken, S.E., and Bezanilla, M. (2023). CRISPR-Cas9 genome editing in the moss *Physcomitrium* (formerly *Physcomitrella*) *patens*. Current Protocols 3, e725. 10.1002/cpz1.725.

111. Wu, S.-Z., and Bezanilla, M. (2014). Myosin VIII associates with microtubule ends and together with actin plays a role in guiding plant cell division. eLife 3, e03498. 10.7554/eLife.03498.

112. Clough, S.J., and Bent, A.F. (1998). Floral dip: a simplified method for *Agrobacterium*-mediated transformation of *Arabidopsis thaliana*. Plant J 16, 735–743. 10.1046/j.1365-313x.1998.00343.x.

113. 113. Niu, Y., and Sheen, J. (2012). Transient expression assays for quantifying signaling output. In Plant Signalling Networks: Methods and Protocols, Z.-Y. Wang and Z. Yang, eds. (Humana Press), pp. 195–206. 10.1007/978-1-61779-809-2_16.

114. Wang, J., Zhang, X., Greene, G.H., Xu, G., and Dong, X. (2022). PABP/purine-rich motif as an initiation module for cap-independent translation in pattern-triggered immunity. Cell 185, 3186–3200.e17. 10.1016/j.cell.2022.06.037.

